# A Minimal Regulatory Spatial Model for Emergent Multicellular Organization in Dissipative Environments

**DOI:** 10.64898/2026.04.24.720740

**Authors:** Luca Zammataro

## Abstract

How multicellular organization, morphology, and functional diversity emerge from minimal regulatory principles remains a central question in theoretical biology and systems medicine. Here we introduce **Evoscope**, a spatial computational framework in which tissue-like organization arises from a compact regulatory grammar governing nutrient uptake, adhesion, motility, competition, protection, and identity commitment. Cells evolve on a toroidal hexagonal lattice within a dissipative environment shaped by nutrient diffusion, energetic constraints, proliferation, death, and local interactions. Despite its minimal rule set, Evoscope reproducibly generates aggregate formation, territorial expansion, competitive interfaces, differentiated colonies, and transient ecological niches. Beyond serving as a minimal model of emergent multicellular organization, Evoscope is designed as a controlled testbed in which the relationship between internal regulatory state and collective morphology is fully accessible. Cell identities are established through three binary commitment variables that define eight heritable cluster states, each associated with distinct balances of cohesion, invasiveness, metabolic efficiency, and competitive fitness. These programs give rise to coherent multicellular colonies with characteristic spatial behaviors, including collective movement, transient persistence, and eventual dissolution.

To test whether visible morphology encodes information about hidden internal dynamics, we trained convolutional autoencoders with supervised prediction heads on simulation snapshots. The resulting latent spaces displayed smooth temporal organization and structured low-dimensional manifolds, indicating that evolving multicellular configurations occupy non-random, learnable regions of state space. Moreover, internal regulatory profiles could be partially inferred from morphology alone, supporting the view that collective form functions as a mesoscopic encoding of underlying regulatory dynamics.

Evoscope therefore provides a proof of concept that autoencoder-based representation learning can recover informative latent structure in a synthetic multicellular system whose internal rules are known. More broadly, these results support the hypothesis that, given sufficiently rich paired morphological and spatial-transcriptional data, related approaches may help identify latent mesoscopic variables in real biological systems.

## 1 Introduction

Understanding how multicellular organization emerges from minimal intracellular regulatory mechanisms remains a central challenge in biology. While molecular biology has provided detailed insights into gene regulation at the molecular level, tissue-level observations describe structure at higher organizational scales, but the intermediate regime in which collective organization arises remains poorly characterized. This intermediate regime, often referred to as the *mesoscopic scale*, is where local interactions, energy constraints, and regulatory dynamics give rise to structured yet dynamic configurations. This view is consistent with the broader tradition of non-equilibrium self-organization, in which structured collective regimes emerge and persist through dissipative processes operating far from equilibrium [1].

Multicellular organization emerges from the interplay of intracellular regulation, local interactions, spatial constraints, and resource availability. The idea that simple local rules can generate highly structured collective behavior has deep roots in cellular automata theory, from von Neumann’s self-reproducing automata to later studies of emergent complexity in rule-based systems [2, 3]. More specifically, the question of how spatial form emerges from local interactions has long been central to theories of morphogenesis, beginning with Turing’s seminal work on reaction-diffusion systems [4]. More recently, a broad family of computational approaches has been developed to study such processes, including lattice-based, vertex-based, cellular Potts, and other individual-based frameworks for tissue self-organization [5].

Established frameworks such as Morpheus have shown that cell-based mechanics, intracellular ODE models, and extracellular reaction–diffusion fields can be integrated within flexible multiscale simulators [6]. A parallel line of work has explicitly connected multicellular spatial models to intracellular regulatory programs. In particular, multiscale frameworks coupling gene regulatory networks to tissue dynamics have shown how intracellular regulation can shape cell-cycle behavior, differentiation, and spatial organization [7]. Related studies on the emergence of multicellularity and proto-organismal organization have further highlighted how stochastic differentiation, adhesion, and local ecological interactions can produce coherent multicellular aggregates from minimal rule sets [8]. More broadly, although the present framework does not explicitly model any specific signaling pathway, it is loosely inspired by biological patterning principles in which short-range cell–cell interactions can break symmetry and generate local fate bias, boundary formation, and multicellular patterning, as classically described for Notch-mediated lateral inhibition and lateral induction [9].

Recent work on neural cellular automata suggests that cellular-automaton-like developmental dynamics may admit compact latent representations, with families of NCA models organized within learned manifold representation [10]. More broadly, variational autoencoder frameworks have been extended to sequential and dynamical data, supporting the use of latent spaces not only for compressing individual simulation snapshots, but also for representing temporal trajectories of evolving systems [11]. Related work has further shown that autoencoders can extract low-dimensional latent descriptions of biological regulatory dynamics, as illustrated in studies of the *Drosophila* gap-gene network [12]. In parallel, recent reviews have highlighted the growing role of generative and representation-learning approaches in morphogenesis and developmental modeling [13]. Finally, other authors have shown that disentangled variational autoencoders and graph-based representation learning can recover biologically meaningful latent structure from single-cell gene-expression data and help model differentiation processes [14]. Together, these studies motivate the possibility of using autoencoder-based latent spaces to represent trajectories of multicellular organization within a mesoscopic state space.

Although these frameworks provide important conceptual and methodological foundations, they are generally not formulated around the specific question addressed here: whether a minimal multicellular system can generate a biologically interpretable mesoscopic level of organization linking heritable intracellular programs to transient colony-level structure. In other words, while prior approaches successfully model tissues, regulatory programs, or latent manifolds of dynamics, they do not explicitly ask whether stable or metastable multicellular colonies can emerge as compact mesoscopic units preserving lineage identity, collective behavior, and dissipative organization.

Here we introduce *Evoscope*, a minimal gene-regulated multicellular simulation designed to test whether a compact set of functional roles is sufficient to generate stable or metastable mesoscopic structures. The conceptual goal is not to reproduce full biochemical detail, but rather to identify a minimal functional grammar from which multicellular order can re-emerge as a biological regime. In this sense, the model also aligns with the broader idea that biological order may emerge from compact rule systems whose collective consequences are not trivially reducible to their individual components [15]. Indeed, *Evoscope* probes a regime in which collective structures are neither reducible to isolated cells nor adequately described as homogeneous tissues, but instead appear as coherent, lineage-bearing, resource-dependent colonies with their own transient dynamics. More specifically, the model asks whether heritable identity states, local adhesion, motility, killing, and nutrient recycling are sufficient to generate emergent multicellular colonies, collective displacement, transient territoriality, and eventual dissolution under energetic constraints. In our framework, cells are represented as agents on a toroidal hexagonal lattice (*TorHex*) and governed by a minimal internal regulatory unit (*genomoid*) encoding a compact functional grammar (Figure 1).

**Figure 1:**
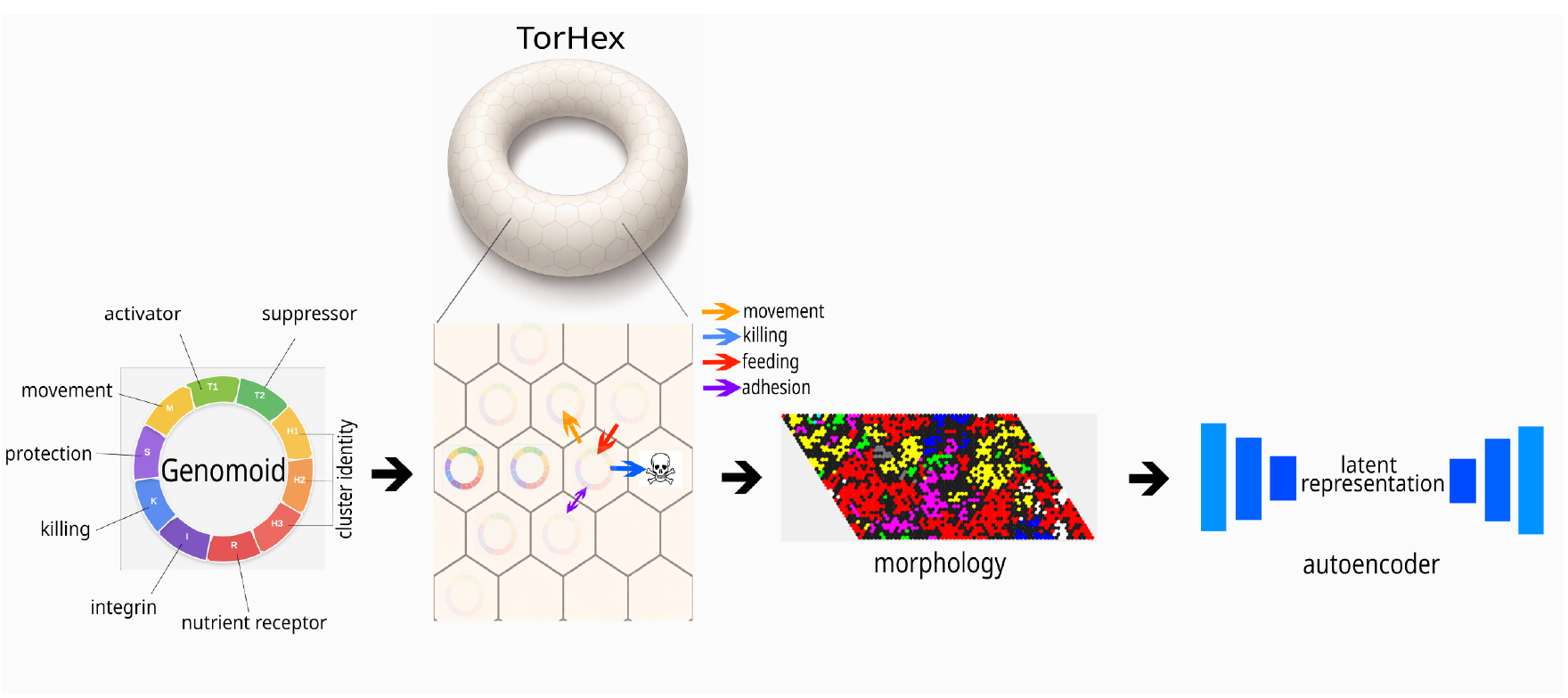
Conceptual overview of the Evoscope framework. Evoscope links three levels of description. On the left, a minimal circular regulatory genome (“genomoid”) defines a compact functional grammar of cellular roles, including activation, repression, movement, adhesion, nutrient uptake, protection, killing, and cluster identity. In the center, these rules are executed by cells on a toroidal hexagonal lattice (TorHex), where local interactions generate differentiation, competition, and emergent colony morphologies. On the right, the resulting spatial organizations are compressed into autoencoder-based latent representations, enabling a mesoscopic description of collective multicellular states.

Cell identities emerge through a heritable commitment process based on three binary histological variables (*H*1, *H*2, and *H*3). The model integrates three essential components: intracellular regulatory dynamics, spatial interactions including adhesion, movement, competition, and local killing, and a shared energetic environment in which nutrients are consumed, redistributed, and recycled through cell death. Through the coupling of these elements, cells compete for space and resources, form clusters based on local recognition rules, and dynamically reorganize their spatial configuration through a balance of cooperative and competitive interactions. A schematic overview of the regulatory architecture and interaction logic is provided in Figure 2. A key feature of the model is that multicellular structure is not imposed, but emerges from the interaction between intracellular state and local environmental conditions. In particular, transcriptional activity is modulated by nutrient availability and local crowding, giving rise to heterogeneous functional states even among genetically identical cells. The model also incorporates polarized and locally resolved behaviors, and spatially resolved competitive expansion.

**Figure 2:**
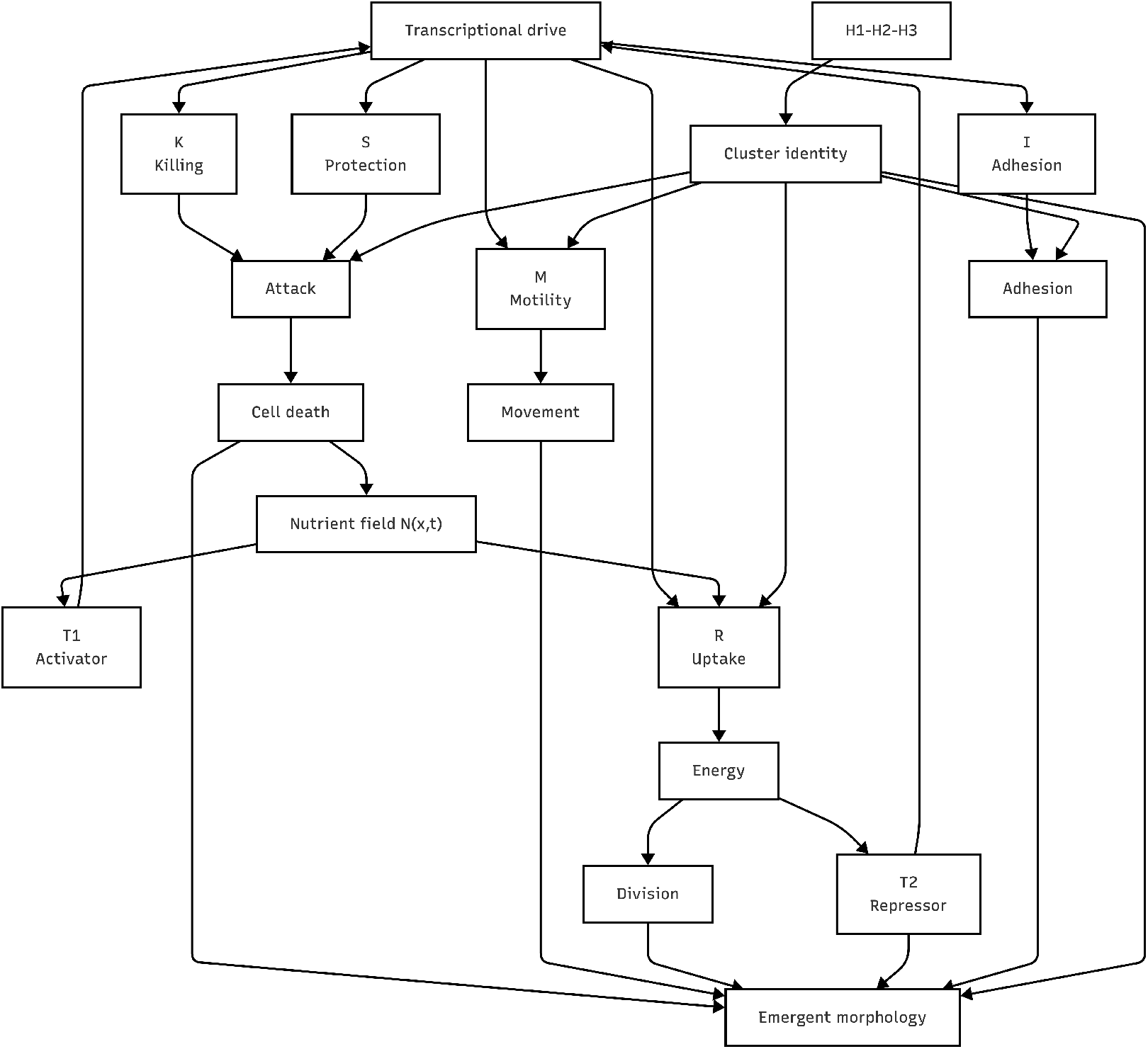
Regulatory architecture of the minimal agent-based genome in *Evoscope*. Schematic representation of the functional interactions linking intracellular regulatory variables, environmental dynamics, and observable cellular behaviors. Nutrient availability feeds the activator module (**T1**) and the uptake program (**R**), which together regulate energy acquisition and support cell-cycle progression and division. The repressor module (**T2**) provides inhibitory feedback within the transcriptional control layer. Adhesion (**I**), motility (**M**), competitive killing (**K**), and protection (**S**) govern local spatial interactions including persistence, movement, attack, and resistance to attack. Binary identity markers (**H1–H3**) define cluster commitment states that modulate collective organization. Cell death returns resources to the nutrient field, closing the energetic recycling loop. The combined action of these components generates emergent multicellular morphologies at the tissue scale.

Because the full internal state of the system is known, *Evoscope* provides a controlled setting to test whether morphology carries recoverable information about hidden regulatory dynamics. In this work, we present the model formulation, characterize its emergent behaviors, and investigate the relationship between spatial organization and underlying regulatory states.

## 2 Methods

The present framework builds on established traditions in agent-based multicellular modeling and latent representation learning [5, 10, 11], while introducing an original minimal integration of regulatory commitment, ecological interaction, and morphology-to-state inference tailored to the mesoscopic question investigated here.

### 2.1 Model Overview

Evoscope is a spatially explicit agent-based model designed to investigate the emergence of organization from minimal intracellular regulatory rules. The system consists of interacting hexagonal cells embedded in a two-dimensional toroidal lattice (*TorHex*) coupled to a diffusive nutrient field. Periodic boundary conditions remove artificial borders by connecting opposite edges, thereby creating a continuous wrap-around geometry equivalent to the topology of a torus.

Each agent is governed by an internal regulatory unit that we term a *genomoid*, a minimal circular architecture encoding a compact set of functional roles. Rather than explicitly reproducing the full molecular cascade of gene expression, the model represents genes through their effective protein concentrations, allowing us to focus on the functional consequences of regulation rather than on detailed biochemical implementation. In this way, the genomoid acts as a minimal interface between intracellular regulatory state, heritable identity, and emergent multicellular phenotype (Figure 1).

Cells evolve in discrete time and interact locally with their six nearest neighbors. The model integrates intracellular regulatory dynamics, energy metabolism, spatial interactions, and stochastic decision-making. At initialization, each agent receives stochastic regulatory states sampled from predefined probability distributions. Continuous activity variables are drawn from bounded uniform distributions, whereas binary identity markers are independently sampled from Bernoulli distributions (Table 1).

**Table 1:** Initial regulatory state of the minimal agent *genomoid*. Protein activities were randomly initialized within bounded ranges to introduce controlled heterogeneity across agents while preserving comparable starting conditions. Directional variables were initialized as six-component vectors whose total abundance was distributed across neighboring directions. Binary identity markers were sampled independently.

| Variable | Functional role | Initialization |
| --- | --- | --- |
| T1 | Activator / nutrient-responsive regulator | Uniform(0.2, 1.2) |
| T2 | Repressor / stress-crowding regulator | Uniform(0.1, 0.8) |
| I | Integrin / Adhesion (directional) | Total Uniform(0.2, 1.0) |
| R | Nutrient uptake / receptor module | Uniform(0.2, 1.0) |
| M | Motility (directional) | Total Uniform(0.2, 1.0) |
| K | Competitive killing (directional) | Total Uniform(0.1, 0.7) |
| S | Surviving / protection / resistance module | Uniform(0.1, 0.7) |
| H1 | Histological Identity bit 1 | Bernoulli(0.5) |
| H2 | Histological Identity bit 2 | Bernoulli(0.5) |
| H3 | Histological Identity bit 3 | Bernoulli(0.5) |

In addition to the intracellular regulatory layer and the multicellular interaction layer, Evoscope includes a shared nutrient environment that contributes to cell-state transitions, competition, survival, and collapse through diffusion, uptake, decay, and release from dead cells. In the present study, however, this environmental field was simulated as part of the generative process but was not included as an explicit target in the autoencoder analyses, which focused on the relationship between morphology and internal regulatory state.

### 2.2 Cellular State Variables

Each cell *i* at time *t* is described by the state vector:

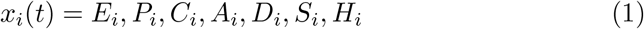

where:

- *E*_*i*_ is the intracellular energy
- *P*_*i*_ = *p*_*i*1_, …, *p*_*i*10_ are protein levels
- *C*_*i*_ is the cell cycle phase (G, S, M, STALL)
- *A*_*i*_ is cellular age
- *D*_*i*_ is the number of divisions
- *S*_*i*_ is the stress state
- *H*_*i*_ is the cluster identity

### 2.3 Cluster Identity and Commitment

Each cell carries a binary identity code:

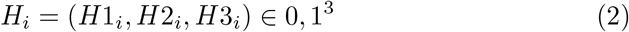

resulting in eight possible committed cluster identities. The binary configuration of the three identity genes (*H*1, *H*2, and *H*3) is converted into an integer cluster label ranging from 0 to 7, corresponding to the eight possible candidate cluster states (2^3^ = 8) (Table 2).

**Table 2:**
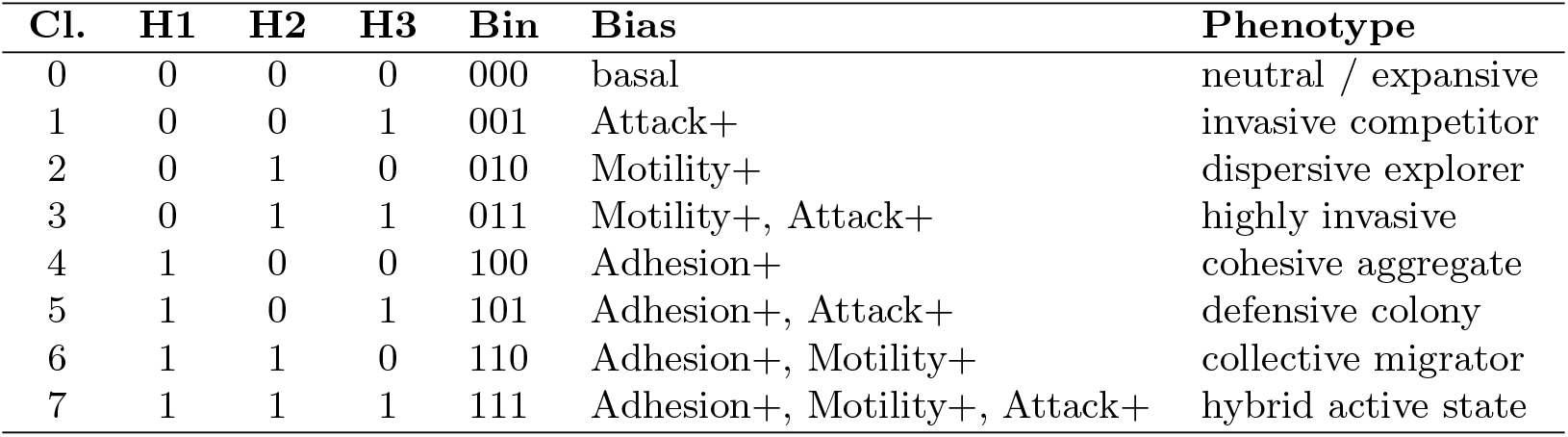
Cluster commitment states defined by the binary identity regulators H1–H3. Stable combinations of the three identity markers generate eight possible committed cluster states. Cells that have not yet stabilized a binary configuration remain in the undetermined state (`cluster_id = None`).

| Cl. | H1 | H2 | H3 | Bin | Bias | Phenotype |
| --- | --- | --- | --- | --- | --- | --- |
| 0 | 0 | 0 | 0 | 000 | basal | neutral / expansive |
| 1 | 0 | 0 | 1 | 001 | Attack+ | invasive competitor |
| 2 | 0 | 1 | 0 | 010 | Motility+ | dispersive explorer |
| 3 | 0 | 1 | 1 | 011 | Motility+, Attack+ | highly invasive |
| 4 | 1 | 0 | 0 | 100 | Adhesion+ | cohesive aggregate |
| 5 | 1 | 0 | 1 | 101 | Adhesion+, Attack+ | defensive colony |
| 6 | 1 | 1 | 0 | 110 | Adhesion+, Motility+ | collective migrator |
| 7 | 1 | 1 | 1 | 111 | Adhesion+, Motility+, Attack+ | hybrid active state |

Cells do not necessarily begin in a committed state, but may remain undetermined (*u*) until a stable identity is established. Commitment occurs only when the same candidate state persists for a predefined number of consecutive epochs (`commitment_stability_epochs`), introducing temporal memory and preventing switching due to transient fluctuations. Once stabilized, committed identities modulate downstream behavioral biases, including adhesion, motility, aggressiveness, and nutrient uptake. Under prolonged energetic stress, cells may lose commitment and return to a decommitted or undetermined state.

### 2.4 Spatial Environment

Cells occupy sites on a two-dimensional hexagonal lattice of size *W* × *H* with periodic (toroidal) boundary conditions. Each lattice site *x* is associated with a scalar nutrient concentration *N* (*x, t*).

Each cell interacts with its six neighboring positions:

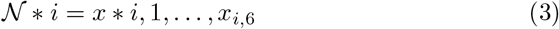

### 2.5 Global Simulation Parameters

The behavior of the system is governed by a set of global simulation parameters controlling initialization, nutrient diffusion, energetic constraints, cell-cycle transitions, regulatory kinetics, commitment dynamics, and spatial decision biases. Unless otherwise stated, default values reported in Table 3 were used throughout the study.

**Table 3:**
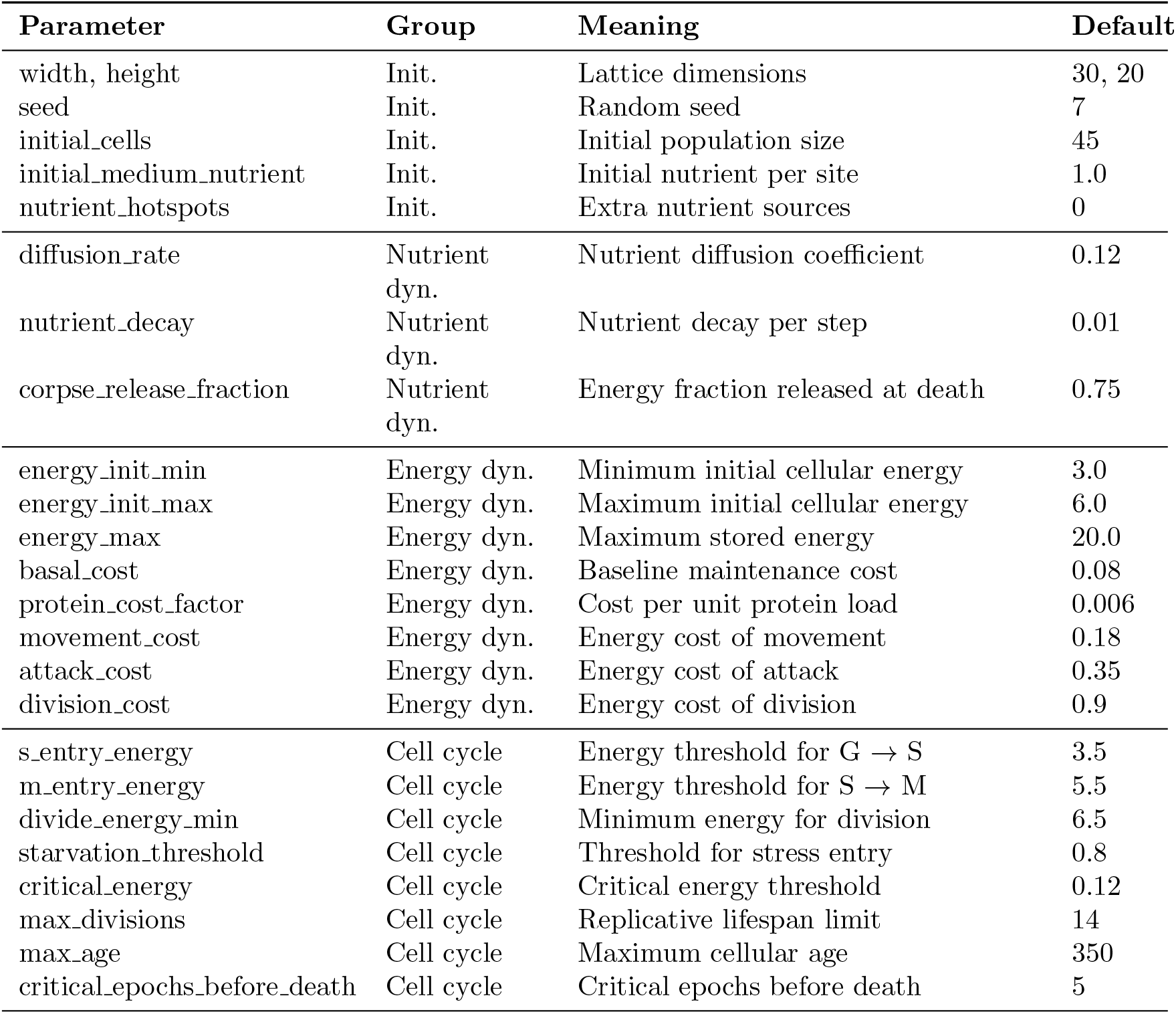
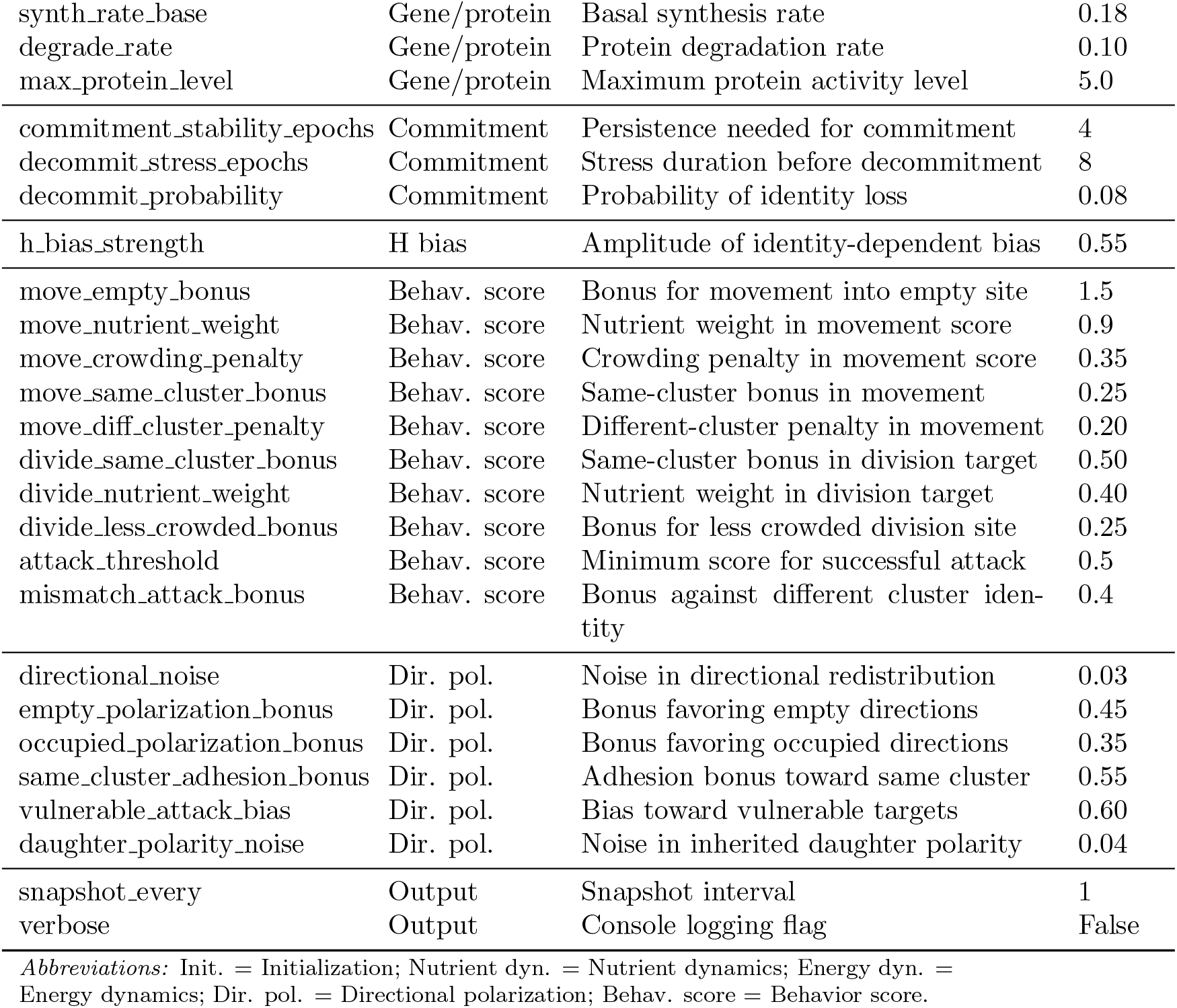
Core global simulation parameters used in Evoscope. Parameters are grouped according to their functional role in the model.

These parameters were selected to generate stable and interpretable dynamical regimes rather than to fit any specific biological dataset. Parameter values were chosen through exploratory simulation to ensure persistence of multicellular dynamics, coexistence of multiple behavioral regimes, and numerical stability across runs.

### 2.6 Gene/Protein Regulatory Dynamics

The model includes the *genomoid*, a structure containing 10 genes encoding proteins, with distinct functional roles: transcriptional regulators (*T* 1, *T* 2), membrane proteins (*I, R, M*), interaction proteins (*K, S*), and identity markers (*H*1, *H*2, *H*3) (Table 1 and Table 2). The minimal regulatory genome implemented in each agent consists of ten functional variables: **T1** and **T2** (transcriptional regulators responsive to nutrient availability, crowding, and stress), **I** (an integrin-like adhesion program controlling cohesion and local persistence), **R** (a receptor/reuptake module governing nutrient acquisition and energetic gain), **M** (a motility program driving directional exploration and spatial expansion), **K** (a killing or competitive effector controlling local displacement of neighboring cells), **S** (a shield/protection module conferring resistance to competitive attack). Together, these variables define a compact functional grammar through which multicellular organization can emerge from local interactions. Initial regulatory values were stochastically sampled from predefined distributions to generate controlled heterogeneity across agents while maintaining comparable starting conditions (Table 1).

#### Transcriptional Regulation

The activities of the transcriptional regulators *T*1_*i*_ and *T*2_*i*_ are updated at each time step as functions of local nutrient availability, local crowding, and stress:

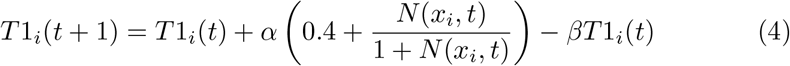

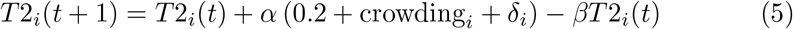

where *x*_*i*_ denotes the lattice position of cell *i, N* (*x*_*i*_, *t*) is the extracellular nutrient concentration at that position and time *t, α* is the basal synthesis coefficient, and *β* is the degradation coefficient. The numerical coefficients in the transcriptional update rules were initially set as heuristic baseline weights to produce a balanced regime between constitutive activation and context-dependent repression. They do not represent fitted biochemical constants, but tunable control parameters of the minimal regulatory grammar.

Local crowding is defined as the fraction of occupied neighboring sites:

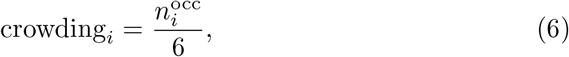

where 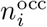 is the number of occupied neighboring sites among the six adjacent lattice positions. The variable *δ*_*i*_ represents stress-dependent activation, which increases when the energetic state of the cell falls below the starvation threshold. Thus, *T*1_*i*_ primarily responds to nutrient availability, whereas *T*2_*i*_ increases with crowding and stress.

#### Global Transcriptional Drive

Gene expression is modulated by a global transcriptional drive that integrates the opposing effects of activation and repression:

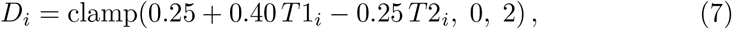

where *D*_*i*_ is the effective transcriptional drive of cell *i, T*1_*i*_ is the activator level, and *T*2_*i*_ is the repressor level. The function clamp(*x, a, b*) truncates the value of *x* to the interval [*a, b*], ensuring that the transcriptional drive remains bounded between 0 and 2. Thus, higher *T*1_*i*_ increases the global drive, whereas higher *T*2_*i*_ reduces it.

#### Protein Dynamics

For the functional effector variables, protein activity is updated through first-order production and degradation dynamics of the general form

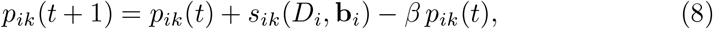

where *p*_*ik*_(*t*) denotes the activity of protein *k* in cell *i* at time *t, D*_*i*_ is the global transcriptional drive, *β* is the degradation coefficient, and *s*_*ik*_(*D*_*i*_, **b**_*i*_) is a protein-specific synthesis term that depends on the transcriptional drive and, when applicable, on identity-dependent bias terms **b**_*i*_. In the present model, the effector proteins *I, R, M, K*, and *S* share this general structure but differ in their synthesis coefficients and bias contributions. By contrast, the identity variables *H*1, *H*2, and *H*3 follow a distinct update rule before commitment, and are subsequently stabilized according to the committed cluster state.

### 2.7 Energy Dynamics

Cellular energy is updated at each time step as the balance between nutrient uptake and metabolic cost:

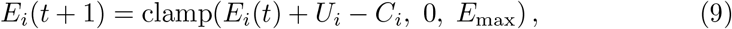

where *E*_*i*_(*t*) is the intracellular energy of cell *i* at time *t, U*_*i*_ is the energy gained through nutrient uptake, *C*_*i*_ is the energetic cost incurred during the same step, and *E*_max_ is the maximum allowed energy level.

Nutrient uptake is defined as

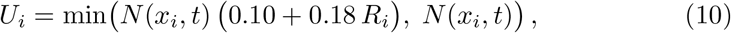

where *x*_*i*_ denotes the lattice position of cell *i, N* (*x*_*i*_, *t*) is the extracellular nutrient concentration at that site, and *R*_*i*_ is the uptake-associated receptor activity. After uptake, the local nutrient field is reduced accordingly.

Metabolic cost is defined as

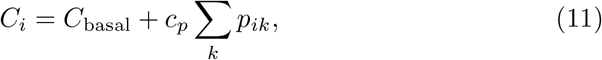

where *C*_basal_ is the baseline maintenance cost, *p*_*ik*_ denotes the level of protein *k* in cell *i*, and *c*_*p*_ is the cost per unit protein load. Additional action-specific costs, including movement, attack, and division, are applied when those actions are executed. Cells enter a stress regime when energy falls below the starvation threshold and undergo death if critically low energy persists for a sufficient number of consecutive steps.

### 2.8 Cell Cycle and Division

Cells transition between cycle phases based on energy thresholds:

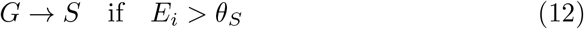

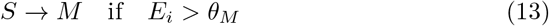

Division occurs when:

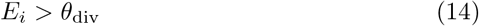

Daughter cells inherit a fraction of the parent’s energy and protein content.

### 2.9 Cell Motility

Cell movement is governed by a local scoring rule evaluated over accessible neighboring lattice sites. Candidate target positions are restricted to empty neighboring sites, and movement is suppressed when adhesive persistence dominates over exploratory drive. In particular, the cell remains stationary when

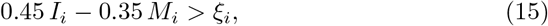

where *I*_*i*_ is the adhesion-related activity, *M*_*i*_ is the motility-related activity, and *ξ*_*i*_ is a random threshold drawn uniformly from the interval [0.1, 1.4].

If movement is allowed, each empty neighboring site *d* is assigned a score

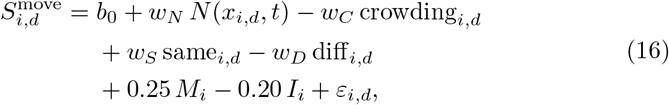

and the selected direction is

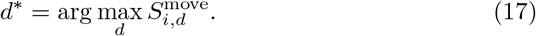

Here, *b*_0_ is the base bonus for moving into an empty site, *N* (*x*_*i,d*_, *t*) is the nutrient concentration at the candidate neighboring position, crowding_*i,d*_ is the local crowding around that site, same_*i,d*_ and diff_*i,d*_ are the numbers of neighboring cells sharing or not sharing the same committed identity, and *ε*_*i,d*_ is a small random perturbation. The coefficients *w*_*N*_, *w*_*C*_, *w*_*S*_, and *w*_*D*_ weight nutrient attraction, crowding penalty, same-cluster attraction, and different-cluster repulsion, respectively. Thus, movement emerges from a balance between exploration, nutrient seeking, local crowding avoidance, and cluster-dependent cohesion. The coefficients *b*_0_, *w*_*N*_, *w*_*C*_, *w*_*S*_, and *w*_*D*_ correspond respectively to `move_empty_bonus, move_nutrient_weight, move_crowding_penalty, move_same_cluster_bonus`, and `move_diff_cluster_penalty` listed in Table 3.

### 2.10 Local Killing and Space Clearing

When a cell in mitotic phase has sufficient energy to divide but no adjacent empty site is available, it may attempt to clear space by attacking a neighboring cell. For each occupied neighboring site *j*, the model computes an attack score

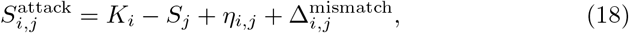

where *K*_*i*_ is the competitive killing activity of the attacking cell, *S*_*j*_ is the protection activity of the neighboring cell, *η*_*i,j*_ is a small random perturbation, and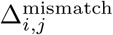 is an additional bonus applied when the two cells belong to different committed cluster identities.

Among all occupied neighbors, the target with the highest attack score is selected. A killing event occurs only if

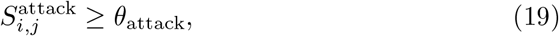

and if the attacker has sufficient energy to pay both the attack cost and the subsequent division cost. Upon a successful attack, the neighboring cell is removed, part of its remaining energy is released back into the local nutrient field, and the attacking cell immediately divides into the cleared site. In this way, local killing acts as a space-clearing mechanism that couples competition to proliferation under crowding.

### 2.11 Nutrient Field Dynamics

The extracellular nutrient field evolves through diffusion, decay, and release from cell death.

#### Diffusion

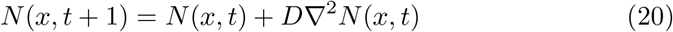

#### Decay

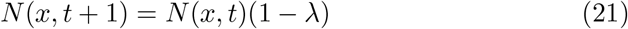

#### Release from Cell Death

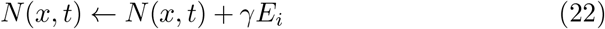

### 2.12 System Dynamics

The system evolves according to:

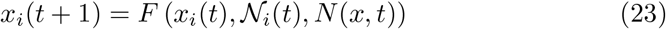

where *F* represents the coupled intracellular, spatial, and environmental dynamics.

### 2.13 Convolutional Autoencoder for Mesoscopic Representation Learning

To investigate whether emergent multicellular morphologies contain recoverable information about internal regulatory states, we trained a convolutional autoencoder coupled to a supervised prediction head. The objective was to learn a low-dimensional latent representation of lattice configurations while preserving both spatial structure and regulatory information.

At simulation step *t*, the spatial configuration of the system was represented as a single-channel lattice image

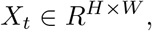

where *H* and *W* denote grid height and width, respectively. Each lattice site encoded the occupancy state of the corresponding hexagonal location, with values representing empty space, undetermined cells, or committed cluster identities. For neural processing, the grid was normalized and treated as a single-channel input image.

The encoder mapped each morphology snapshot *X*_*t*_ to a latent vector

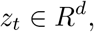

with *d* = 8, using successive convolutional layers followed by a fully connected projection. A decoder reconstructed the original morphology from the latent state using a symmetric transposed-convolution architecture. In parallel, the same latent vector was used by a supervised prediction head to estimate either global or cluster-resolved regulatory variables.

In the global setting, the target vector contained the mean expression of the seven functional variables

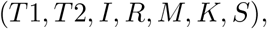

whereas in the cluster-resolved setting the target consisted of the corresponding variables stratified across the eight cluster identities. Training minimized a multitask objective combining morphology reconstruction loss and gene-prediction loss:

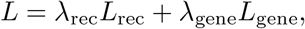

where *L*_rec_ measured reconstruction error and *L*_gene_ measured prediction error on the associated regulatory targets.

The learned latent vector

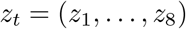

was interpreted as a mesoscopic state of the system, i.e., a compressed representation intermediate between microscopic cell-level variables and macroscopic morphology. Temporal trajectories in latent space were analyzed to identify smooth developmental paths, reorganizations, and attractor-like multicellular regimes.

The simulation framework was implemented in Python, and the autoencoder was implemented in PyTorch [16]. Optimization was performed using Adam with mini-batch gradient descent. Latent vectors, reconstruction losses, and predicted regulatory profiles were exported for downstream analysis and visualization.

## 3 Results

### 3.1 Emergent Spatial Morphogenesis and Population Collapse

Starting from a sparse distribution of cells on the TorHex lattice, the system reproducibly generated a sequence of large-scale spatial states; Figure 3 shows one representative simulation run (seed 42). In the earliest phase (step 5), the population remained fragmented, with isolated cells, small colonies, and several undetermined agents that had not yet stabilized a commitment identity (gray cells in the “u” state; Figure 3A).

**Figure 3:**
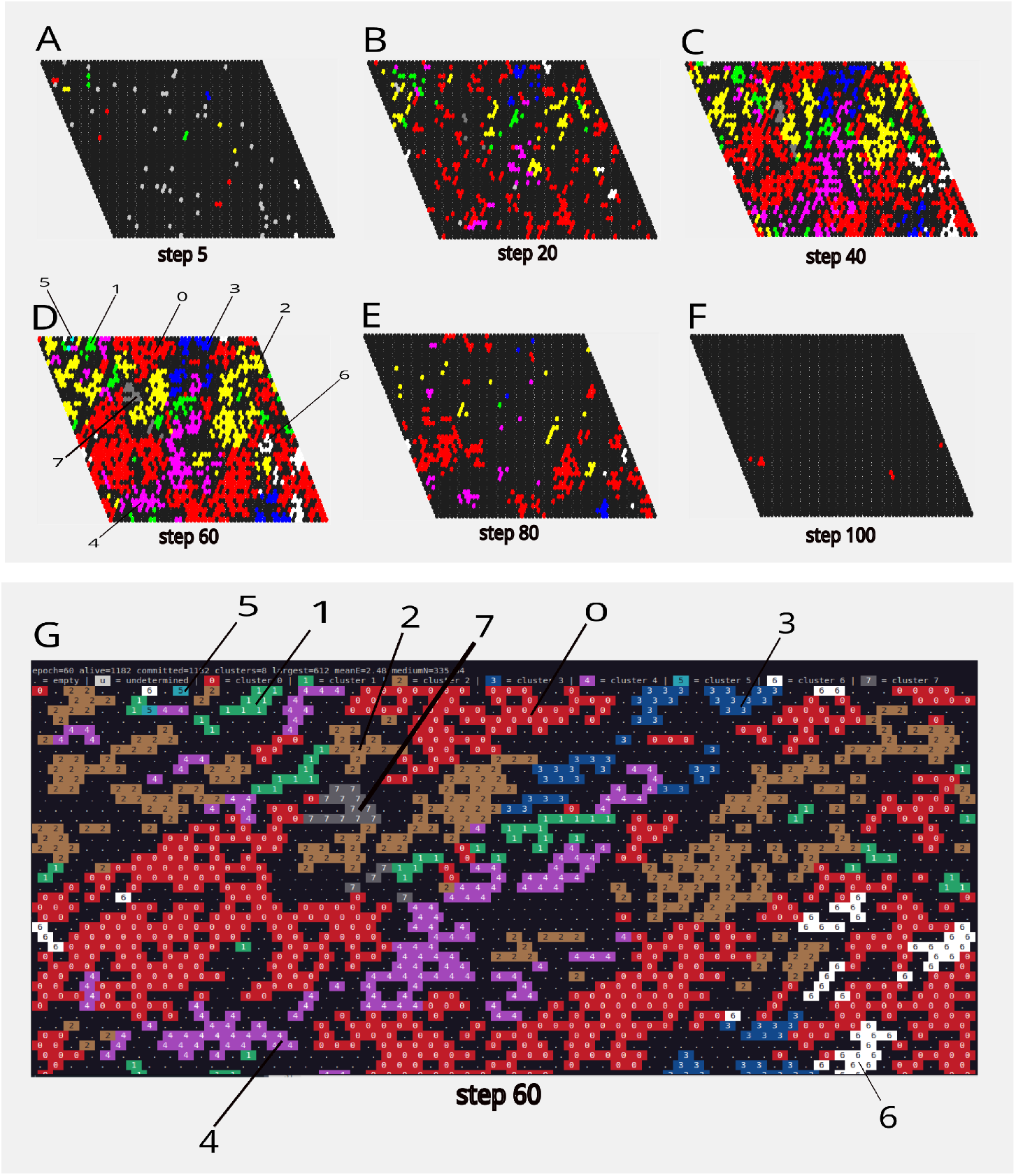
Emergent multicellular morphogenesis in a representative *Evoscope* simulation. Sequential snapshots of the toroidal hexagonal lattice generated under the initial parameter set `--width 60 --height 40 --seed 42 --epochs 150` illustrate the spontaneous formation, diversification, and collapse of multicellular organization from an initially sparse population. Colored hexagons represent living cells committed to stable cluster identities, whereas dark sites indicate empty lattice positions. Gray cells correspond to undetermined or not-yet-committed cells that have not stabilized a binary identity state. (A) Step 5: early exploratory phase dominated by sparse cells, including several undifferentiated gray cells and small nascent colonies. (B) Step 20: rapid population expansion with increasing commitment and local cluster nucleation. (C) Step 40: dense multicellular regime characterized by coexistence of multiple cluster identities and extensive spatial competition. (D) Step 60: peak structural complexity with heterogeneous domains, crowding, and dynamic interfaces between competing populations. Numeric labels indicate representative cluster identities (0–7) that have reached visually stable and recognizable spatial domains. (E) Step 80: collapse phase marked by population decline, fragmentation, and loss of multicellular structure following progressive resource exhaustion. (F) Step 100: near-extinction state in which only a few viable cells remain. (G) Alternative ASCII rendering of the Step 60 configuration. This symbolic representation enhances the visual discrimination of cluster boundaries and identities by explicitly labeling cells according to their committed state, thereby providing a clearer view of spatial domain organization than the high-resolution graphical rendering alone. These simulations show that complex tissue-like spatial organization can emerge transiently from minimal local regulatory rules operating in a dissipative environment.

By step 20, the system underwent rapid population expansion accompanied by the emergence of multiple committed clusters. Distinct local populations became spatially recognizable, and early nucleation events were visible as groups of cells sharing the same identity began to aggregate and expand into multicellular domains (Figure 3B).

At intermediate times (steps 40–60), the lattice reached its highest degree of structural complexity. Multiple cluster identities coexisted across the domain, generating heterogeneous spatial territories separated by dynamic interfaces (Figure 3C–D). During this regime, some populations formed compact and cohesive regions, whereas others appeared more dispersed or locally invasive. Competition for space became especially evident, with boundaries continuously reshaped by neighboring populations.

The late dynamics were characterized by progressive destabilization of the multicellular structure. By step 80, fragmentation and population decline became apparent, and previously coherent domains began to disassemble (Figure 3E). Spatial order was progressively lost as cluster boundaries collapsed and surviving cells became increasingly sparse.

Finally, by step 100, the system approached a near-extinction regime in which only a few viable cells remained (Figure 3F). The overall trajectory therefore consisted of emergence, diversification, peak structural organization, and eventual collapse of multicellular order under fixed initial conditions.

### 3.2 Regulatory Specialization Underlying Emergent Morphologies

To characterize the internal dynamics underlying the spatial states shown in Figure 3, we analyzed the temporal trajectories of the regulatory variables at both global and cluster-resolved levels. As shown in Figure 4, aggregation across 100 independent simulations preserves both the global temporal structure of the system and the cluster-specific regulatory programs, indicating that these dynamics are robust features of the model rather than artifacts of individual runs.

**Figure 4:**
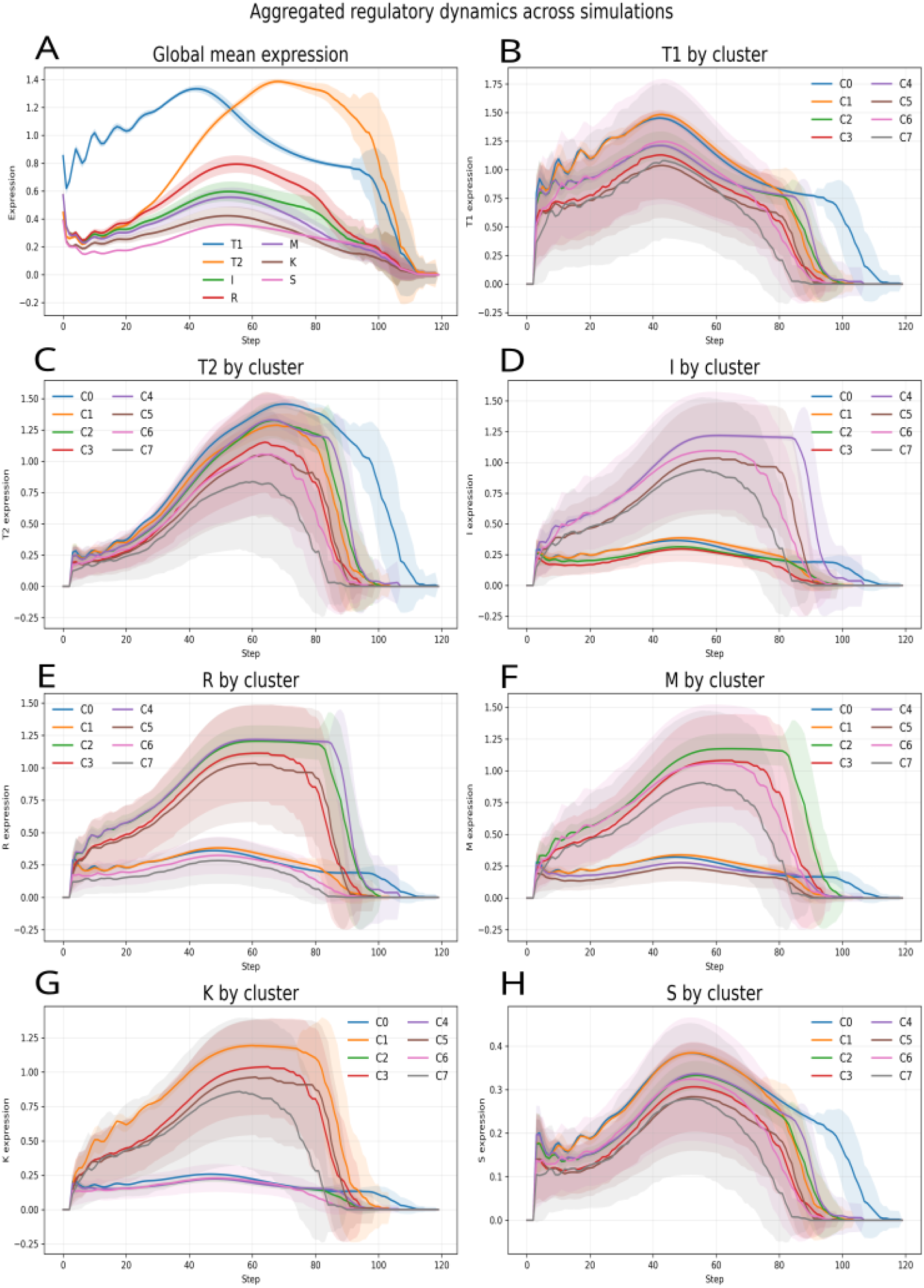
Aggregated temporal dynamics of regulatory variables across 100 independent Evoscope simulations. All simulations were performed using identical global parameters (Table 3), a toroidal hexagonal grid of size 60 × 40, and a duration of 150 steps. Simulations were repeated across 100 independent random seeds (25–124). Solid lines represent the mean expression trajectory across simulations, and shaded regions indicate ±1 standard deviation. (A) Global mean expression of the seven functional protein variables (**T1, T2, I, R, M, K, S**), averaged across all living cells at each step. The aggregated dynamics reveal a reproducible temporal program characterized by early activation, intermediate specialization, and late collapse associated with population extinction.(B–H) Cluster-resolved mean expression trajectories for the same regulatory variables, showing the emergence of distinct functional programs among committed identities. (B) activator module (**T1**); (C) repressor module (**T2**); (D) adhesion program (**I**); (E) nutrient uptake/receptor program (**R**); (F) motility program (**M**); (G) competitive killing program (**K**); (H) protection module (**S**). Different cluster identities reproducibly adopt distinct regulatory strategies, including aggressive high-**K** states, adhesive high-**I** states, motile high-**M** states, and metabolically active high-**R**/**T1** states. The preservation of these patterns across 100 simulations indicates that cluster-specific regulatory programs are robust features of the model rather than artifacts of individual runs.

At the global level (Figure 4A), the system exhibited structured temporal phases rather than random fluctuations. The activator module (**T1**) rose rapidly during the early expansion phase, whereas the repressor module (**T2**) accumulated later and became more prominent under crowded conditions. Additional variables, including adhesion (**I**), motility (**M**), competitive killing (**K**), protection (**S**), and nutrient acquisition (**R**), also followed distinct non-monotonic trajectories over time.

Cluster-resolved profiles (Figure 4B–H) revealed clear diversification among committed identities. Although all clusters emerged from the same underlying rule set, they occupied distinct regulatory states over time. Some clusters were characterized by comparatively higher nutrient uptake (**R**), others by elevated killing activity (**K**), whereas additional populations showed increased adhesion (**I**) or motility (**M**). Protective activity (**S**) also varied across populations over time.

These differences were accompanied by distinct temporal behaviors, including early expansion, sustained persistence, transient dominance, and late decline depending on cluster identity. Several trajectories converged toward simultaneous reduction during the final stages of the simulation, consistent with the population collapse observed in Figure 3E–F.

These cluster-specific programs emerged from the binary identity markers **H1**–**H3**, which define eight possible commitment states. Multiple states were simultaneously represented during the multicellular phase of the dynamics.

Overall, these results indicate that a minimal regulatory system can generate phenotypic heterogeneity, robust diversification into multiple coexisting populations, and dynamically evolving multicellular organization under fixed environmental conditions.

### 3.3 Multitask autoencoder analysis with global targets

A multitask convolutional autoencoder was trained using morphology snapshots as input and global regulatory variables as associated targets. The same latent representation was used for snapshot reconstruction and target prediction.

Predicted trajectories for multiple global variables followed the corresponding simulated temporal profiles across the dataset. Reconstruction and prediction losses decreased during training and converged to stable values.

The latent coordinates extracted from the trained model showed ordered temporal trajectories across simulation time (Figure 5A). Several variables changed progressively during the simulation, whereas others remained within narrower ranges. In the two-dimensional projection defined by (*z*_1_, *z*_2_), consecutive time points formed a continuous trajectory through latent space (Figure 5C).

**Figure 5:**
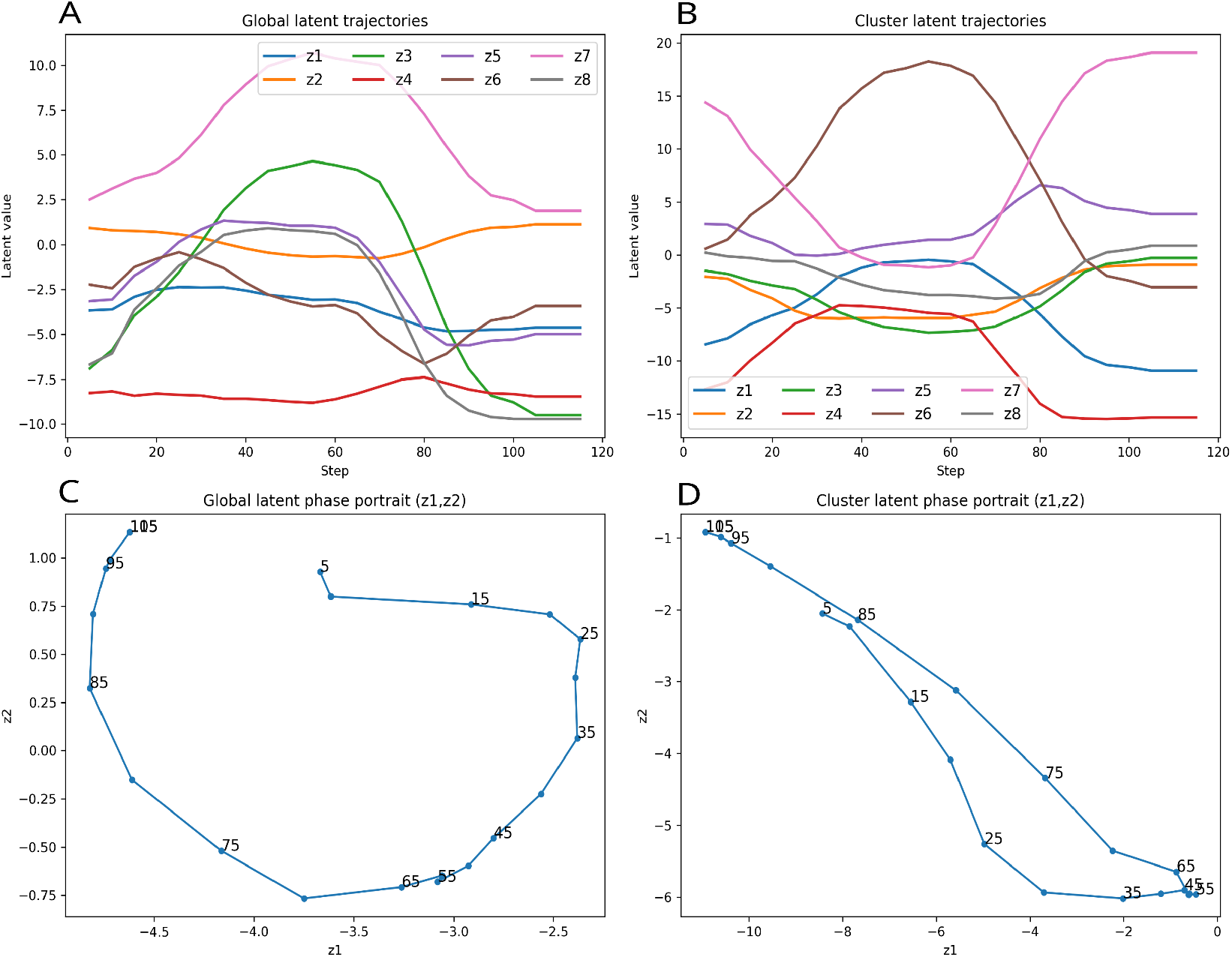
Temporal organization of the learned latent mesoscopic spaces. All panels correspond to the simulation generated under the reference parameter set `--width 60 --height 40 --seed 42 --epochs 150`. (A) Temporal trajectories of the eight latent variables learned by the multitask autoencoder when trained against global regulatory targets. The latent coordinates evolve smoothly over simulation time and exhibit coordinated changes rather than random fluctuations, indicating that the global latent space captures structured system-level transitions. (B) Temporal trajectories of the eight latent variables learned when training against cluster-resolved targets. In this setting, the latent coordinates display sharper divergence and stronger heterogeneity, consistent with the emergence of cluster-specific ecological and functional regimes. (C) Two-dimensional phase portrait of the global latent trajectory in the (*z*_1_, *z*_2_) plane, with labels indicating simulation steps. The trajectory follows a coherent path through latent space, suggesting that the evolving multicellular system occupies an organized low-dimensional manifold rather than an unstructured cloud of states. (D) Two-dimensional phase portrait of the cluster-resolved latent trajectory in the (*z*_1_, *z*_2_) plane. Compared with the global case, the cluster-level trajectory spans a broader and more asymmetric region of latent space, reflecting the stronger contribution of internal population structure and differentiated cluster dynamics. Together, these panels support the interpretation of the learned latent coordinates as mesoscopic state variables linking regulatory dynamics to emergent multicellular morphology.

To further interpret the learned representation, correlations were computed between latent coordinates and observed regulatory variables (Figure 6A). Several dimensions showed broad positive associations with multiple factors, indicating that the encoder captured coordinated system-wide states rather than isolated variables. In particular, *z*_3_, *z*_4_, and *z*_8_ were strongly correlated with adhesion, uptake, motility, aggressiveness, and stress-related programs, whereas *z*_2_ and *z*_6_ displayed negative associations with the repressive factor *T* 2.

**Figure 6:**
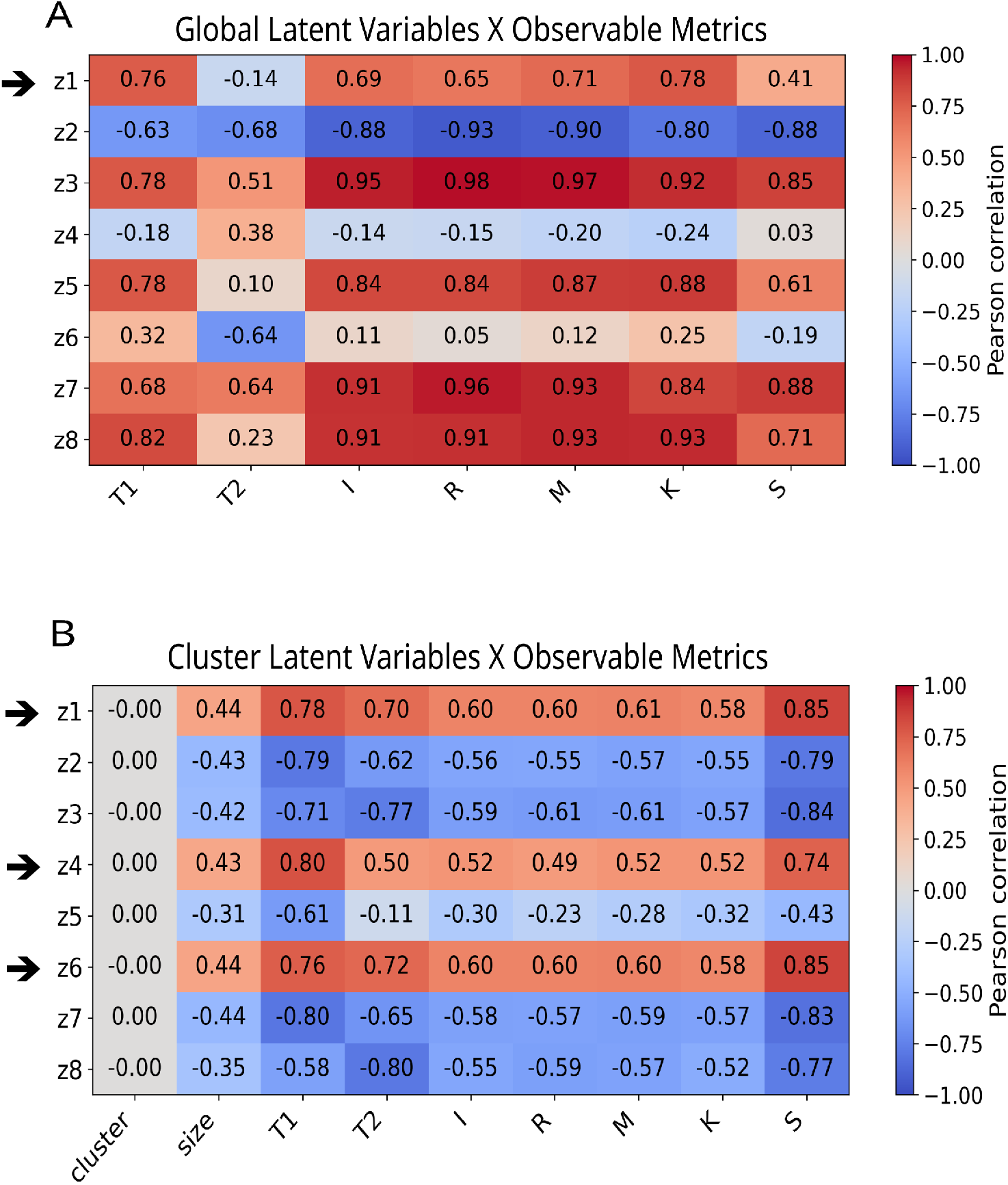
Correlation structure of the learned latent spaces. (A) Heatmap of Pearson correlations between the global latent coordinates and observable variables derived from system-wide averages, shown for the simulation generated under the reference parameter set `--width 60 --height 40 --seed 42 --epochs 150`. Several latent dimensions show strong positive associations with activating and operational variables (*T*1, *I, R, M, K*), whereas others are more closely associated with repressive or stabilizing components (*T*2, *S*). This indicates that different coordinates capture distinct functional programs or global system states. (B) Heatmap of Pearson correlations between cluster-resolved latent coordinates and clusterlevel observables. Some dimensions are positively associated with cluster size and multiple functional variables, consistent with persistent and highly active population states, whereas others show broadly negative correlations, indicative of declining or low-fitness regimes. The negligible correlation with numerical cluster identity indicates that the latent representation encodes ecological and functional states rather than arbitrary labels. Together, these analyses support the interpretation of latent variables as mesoscopic coordinates linking microscopic regulatory factors to emergent multicellular organization.

### 3.4 Multitask autoencoder analysis with cluster-resolved targets

The same computational architecture was next trained using cluster-resolved regulatory states as targets instead of global averages, while preserving the same reconstruction objective and latent dimensionality.

Predicted cluster-level outputs retained temporal correspondence with the simulated data, and training losses converged during optimization.

The latent coordinates obtained in this setting also followed ordered temporal trajectories (Figure 5B). Compared with the global-target setting, several latent variables spanned wider numerical ranges and exhibited sharper transitions during intermediate simulation stages.

In the corresponding (*z*_1_, *z*_2_) projection, consecutive time points again formed a continuous path through latent space (Figure 5D), with a geometry distinct from that observed for global targets.

Correlation analysis of the cluster-resolved latent variables revealed a sharper and more heterogeneous structure (Figure 6B). Some dimensions were positively associated with cluster size and activation-related factors, whereas others showed inverse relationships with the same variables. This pattern is consistent with latent coordinates that encode internal population heterogeneity and cluster-specific organizational states beyond the information contained in global averages.

## 4 Discussion

### 4.1 Minimal Regulatory Programs as Drivers of Multicellular Patterning

The present study introduces a minimal computational framework in which multicellular organization emerges from the interaction of a small number of interpretable regulatory variables embedded in a dissipative spatial environment. Although the model is intentionally simplified, the resulting behaviors are non-trivial: cells self-organize into structured aggregates, diversify into distinct populations, compete for space and resources, and generate morphology that retains measurable relationships with hidden internal states. Taken together, these findings support the broader idea that complex collective organization does not necessarily require highly detailed molecular descriptions or large regulatory networks, but can arise from compact and context-dependent rule systems.

A central result of the simulations is the spontaneous appearance of ordered spatial regimes from initially sparse and weakly structured initial conditions. The progression from exploratory single cells to nucleated clusters, followed by coexistence of heterogeneous domains and eventual collapse, suggests that multicellular organization can be understood as a transient dynamical process rather than as a fixed endpoint. In this sense, tissue-like form is not imposed externally but assembled through iterative local interactions among growth, movement, adhesion, competition, and resource use. The emergence of recognizable developmental phases from minimal assumptions reinforces the view that large-scale biological order may often reflect generic dynamical principles. The early transition toward organized multicellular domains appears to coincide with a specific regulatory regime in which the activating variable **T1** becomes dominant while the repressive variable **T2**, remains comparatively low (Figure 4A-C-D). Under these conditions, local populations stabilize into distinct commitment states and act as nucleation centers for subsequent cluster expansion. This suggests that the onset of large-scale structure may depend on transient asymmetries between positive and negative regulatory drives rather than on any externally imposed organizer.

The temporal sequence observed in the model also highlights the importance of ecological constraints. Population decline was not triggered by an external perturbation or by the failure of a single regulatory component. Instead, collapse emerged endogenously from the coupled consequences of successful expansion: consumption of shared resources, increasing crowding, competitive turnover, and imperfect recycling. This behavior resembles a systems-level carrying-capacity effect in which growth creates the conditions for its own limitation. More generally, the results emphasize that persistence of multicellular organization depends not only on mechanisms of proliferation and differentiation, but also on sustainable coupling between metabolism and environment.

A particularly informative aspect of the model is the explicit association between internal regulatory programs and emergent spatial morphology. Populations characterized by different regulatory balances generated distinct spatial phenotypes across the TorHex lattice. In particular, clusters 4 and 6 formed compact and cohesive domains, a behavior consistent with their elevated expression of the variable **I** (adhesion-related; see Figure 4G), whose role in the model is to promote local aggregation and territorial cohesion. By contrast, clusters 1 and 3 more frequently displayed dispersed or locally invasive configurations together with higher values of the competitive variable **K** (killing; Figure 4E), suggesting that aggressive boundary dynamics can emerge from increased offensive capacity.

An additional nuance emerges from the comparison between clusters 1 and 3. Although both populations displayed elevated values of the competitive variable **K**, they did not share the same profile for the motility-associated factor **M** (Figure 4H). Cluster 3 exhibited higher **M** activity, whereas cluster 1 remained comparatively lower in this dimension. This suggests that similar offensive potential does not necessarily translate into identical spatial behavior. Rather, competitive capacity appears to be modulated by the broader regulatory context: high killing combined with increased motility may favor more dynamic or invasive expansion, whereas high killing in a less motile population may produce a more localized and territorially constrained aggressive phenotype. More generally, the results indicate that emergent morphology depends on multidimensional regulatory balance rather than on any single variable taken in isolation.

An interesting emergent feature of the simulations is that highly aggressive populations were not necessarily the most successful in the long term. Clusters exhibiting elevated killing activity often achieved strong local dominance, yet they did not always produce the broadest territorial expansion or longest persistence. By contrast, less aggressive populations frequently occupied larger regions and survived longer. This apparent paradox is consistent with a classical ecological trade-off between competitive intensity and energetic sustainability. Aggressive strategies may provide immediate territorial advantages, but they also increase turnover, impose energetic costs, and accelerate local resource depletion. More moderate strategies can instead allocate resources toward growth, maintenance, and stable colonization. Even in this minimal artificial ecology, local winners were not always global winners.

More generally, populations dominated by stronger adhesion-like activity tended to stabilize compact territories, whereas populations with elevated competitive or motility-associated activity more often generated dispersed fronts, invasive borders, or unstable territorial interfaces. This interpretation is also qualitatively supported by the presence of local “cavities” or zones of reduced occupancy surrounding the more aggressive clusters, (Figure 3G), consistent with intensified conflict and local population turnover at their boundaries.

These correspondences indicate that visible spatial organization can act as an external signature of hidden internal state. In real biology, the molecular details are vastly richer than in the present model, but the principle may be similar: collective form is shaped by the balance of underlying functional programs, and therefore carries partial information about them.

### 4.2 Latent Mesoscopic Spaces for Tissue Interpretation and Design

These considerations naturally extend the present framework toward emerging areas such as digital pathology and spatial transcriptomics, where tissue architecture is increasingly interpreted not merely as descriptive morphology, but as a proxy for underlying biological state [17]. In Evoscope, the full internal state of the system is known by construction, making it possible to test directly whether morphology alone retains recoverable information about hidden regulatory dynamics. The ability of the multitask autoencoder to predict components of regulatory state from morphology supports this premise.

The learned latent coordinates reinforce this view. Rather than evolving as random scatter, they followed coherent temporal trajectories across simulation time, consistent with the existence of low-dimensional mesoscopic regimes linking microscopic regulatory processes to higher-order spatial organization. Some coordinates varied smoothly, whereas others changed more abruptly during periods of reorganization, suggesting that multicellular dynamics can be compressed into a compact state space while preserving temporal structure.

The correlation analysis provides a more specific interpretation of these latent dimensions. In the global-target setting (Figure 6A), some coordinates appeared to capture selective functional programs, whereas others reflected broader system-wide activity states. In particular, *z*_1_ was positively associated with activating and operational variables of the model (*T*1, *I, R, M*, and *K*), while showing weaker relationships with the more passive or stabilizing components *T*2 and *S*. This pattern is consistent with an expansion-oriented regime in which growth, movement, adhesion, nutrient uptake, and competitive behavior are jointly enhanced. By contrast, coordinates such as *z*_8_ displayed strong but less selective correlations across multiple observables, suggesting that they encode a more global systems-level activity state rather than a single specialized process.

A complementary picture emerged from the cluster-resolved latent representation (Figure 6B). In this setting, some coordinates, particularly *z*_1_, *z*_4_, and *z*_6_, were positively associated with cluster size and multiple functional variables, consistent with persistent and highly active population states. Other dimensions, including *z*_2_, *z*_3_, *z*_7_, and *z*_8_, showed broadly negative correlations, indicating declining, low-fitness, or exhausted cluster regimes. Notably, numerical cluster identity itself showed negligible correlation with all latent coordinates, indicating that the learned representation does not encode arbitrary labels, but rather ecological and functional states such as persistence, activation, and collapse.

This interpretation remained qualitatively stable across independent simulation seeds, although the individual latent axes were not uniquely aligned from run to run. Such behavior is expected in representation learning, where equivalent internal structure may be expressed through rotated, permuted, or redistributed latent coordinates. What remained reproducible was therefore not the identity of any single axis, but the broader latent–observable correlation structure. Representative pairs of global and cluster heatmaps from multiple seeds are provided in the Supplementary Information and accompanying zip archive, where the consistency of these mesoscopic relationships can be examined in greater detail.

The comparison between global-target and cluster-resolved training is therefore particularly informative. When optimized against global averages, the latent space emphasizes coarse system-level transitions such as expansion, crowding, and decline. When trained against cluster-resolved targets, the representation becomes sharper and more heterogeneous, reflecting the contribution of internal population structure and cluster-specific ecological reorganization. The two settings thus provide complementary views of the same evolving system: one highlights the architecture of the population as a whole, whereas the other reveals its organized internal heterogeneity.

Together, these results support the interpretation of latent variables as mesoscopic coordinates of multicellular organization. In this sense, the latent space does not merely compress morphology, but provides a compact and partially interpretable representation of collective biological state that may prove useful for tissue interpretation, morphogenetic analysis, and future inverse-design strategies.

## 5 Conclusions and Future Directions

This study supports the view that multicellular form is a genuinely mesoscopic phenomenon. Between intracellular mechanisms and organism-level anatomy lies an intermediate regime in which repeated local interactions give rise to stable, interpretable, and dynamically evolving collective structures. In the present framework, this level is visible both in the emergent spatial organization of the simulation and in the learned latent representations extracted from it. Morphology is therefore not merely an output of microscopic rules, but a measurable state variable defined at its own descriptive scale.

Beyond its conceptual implications, the framework also provides a useful experimental sandbox. Because real biological systems rarely offer complete access to all relevant variables, rigorous validation of image-to-state inference methods remains difficult. Here, by contrast, the full generative process is observable. This makes it possible to benchmark machine-learning approaches under controlled conditions while preserving non-trivial emergent organization. In this sense, simplified synthetic systems can serve as valuable intermediate testbeds between abstract theory and experimental biology, in line with the broader artificial-life perspective that seeks to understand living organization by synthesizing its essential properties in controlled artificial systems [18]. From this perspective, Evoscope can also be viewed as a minimal artificial-life sandbox in which emergent multicellular organization can be explored under fully observable conditions [19].

Several limitations should be acknowledged. The regulatory variables are abstract functional modules rather than specific genes or biochemical pathways. The simulations are performed on a two-dimensional lattice with simplified local neighborhoods, whereas real tissues are three-dimensional, mechanically deformable, and shaped by long-range signaling. In addition, parameter values were selected to explore qualitative behavior rather than calibrated against experimental measurements. Accordingly, the framework should not be interpreted as a model of any specific organism or tissue, but rather as a tool for identifying general organizational principles under minimal assumptions.

These limitations also define clear directions for future work. Natural extensions include continuous signaling fields, extracellular matrix effects, mechanical forces, and fully three-dimensional environments. At the representation-learning level, richer architectures may help disentangle causal factors, identify transitions between metastable states, and learn control policies for steering the system toward desired morphologies. A particularly important next step will be to test whether learned mesoscopic latent states can function not only as descriptive encodings of morphology, but also as predictive state variables for future form. In such a setting, regulatory profiles observed at a given time point could be mapped into a latent representation, evolved forward in time, and decoded into future tissue configurations, moving the framework from descriptive mesoscopic compression toward predictive mesoscopic dynamics.

A further limitation is that the shared nutrient environment, although explicitly simulated and dynamically coupled to cell behavior, was not included in the representation-learning analyses. Future work could test whether environmental fields also contribute to the latent mesoscopic structure learned from the system.

A related and especially promising direction is inverse morphogenesis: rather than inferring hidden state from observed morphology, one may seek the inverse mapping from target morphology to regulatory programs capable of generating it. In practical terms, this means asking which combinations of local rules, internal states, or environmental conditions are sufficient to produce a desired multicellular architecture. Such an approach could eventually inform synthetic morphogenesis, programmable tissue engineering, and AI-guided biofabrication. More broadly, the study suggests that useful biological insight may emerge not only from increasing realism, but also from carefully chosen simplification. Minimal models can function as conceptual microscopes, isolating principles that remain difficult to detect in the full complexity of living systems. In this sense, the broader implication is that between genes and tissues there exists a computationally tractable intermediate layer, and that this layer may be measurable, learnable, predictive, and ultimately designable.

## Code Availability

The Evoscope source code is publicly available at: https://github.com/lunanfoldomics/evoscope

## Acknowledgements

The author is grateful to Leonardo Mirandola for recognizing the conceptual parallel between lateral patterning mechanisms described in the Notch literature and the local interaction logic underlying the Evoscope model.

