## Supplementary material for "A Minimal Regulatory Spatial Model for Emergent Multicellular Organization in Dissipative Environments": README_seed_42.txt

Evoscope supplementary archive - seed_42
========================================

This archive contains the simulation outputs and autoencoder-derived files
associated with the representative Evoscope run performed with simulation seed 42,
as referenced in the manuscript:

Luca Zammataro.
A Minimal Regulatory Spatial Model for Emergent Multicellular Organization in Dissipative Environments.

Purpose of this archive
-----------------------
This directory provides the main files used to inspect the morphology,
regulatory trajectories, and latent-space analyses for the representative
simulation seed 42 shown and discussed in the manuscript.

Important note
--------------
The simulation seed (42) determines the underlying Evoscope run.
However, the autoencoder analyses include an additional source of stochasticity
due to random initialization and optimization during neural-network training.
Therefore, exact numerical values in latent coordinates and predictions may vary
slightly across independent autoencoder training sessions, even when using the
same simulation seed.

For this reason, the manuscript emphasizes the qualitative consistency of the
latent-observable relationships rather than exact numerical identity of
individual latent dimensions across runs.

Directory contents
------------------

1. Simulation-level files
   - global_genes.csv
       Global mean trajectories of the regulatory variables across simulation time.
   - cluster_genes.csv
       Cluster-resolved mean trajectories of the regulatory variables across simulation time.
   - evoscope_ascii_frames.txt
       ASCII rendering of simulation states, useful for directly inspecting
       cluster identities and spatial organization.
   - snapshots/
       Numpy arrays containing saved lattice states at successive simulation steps.

2. Global autoencoder outputs
   - global_ae_outputs/latents.csv
       Learned latent coordinates from the global-gene autoencoder workflow.
   - global_ae_outputs/predictions.csv
       Predicted global regulatory variables from the global-gene autoencoder.
   - global_ae_outputs/training_history.csv
       Training history for the global-gene autoencoder.
   - global_ae_outputs/torex_autoencoder.pt
       Saved PyTorch model checkpoint for the global-gene autoencoder.

3. Cluster autoencoder outputs
   - cluster_ae_outputs/latents.csv
       Learned latent coordinates from the cluster-gene autoencoder workflow.
   - cluster_ae_outputs/predictions.csv
       Predicted cluster-resolved regulatory variables from the cluster-gene autoencoder.
   - cluster_ae_outputs/training_history.csv
       Training history for the cluster-gene autoencoder.
   - cluster_ae_outputs/torex_autoencoder.pt
       Saved PyTorch model checkpoint for the cluster-gene autoencoder.

4. Derived visual outputs
   - seed_42_global_latent_heatmap.png
       Heatmap summarizing correlations between global latent coordinates
       and global observable regulatory variables.
   - seed_42_cluster_latent_heatmap.png
       Heatmap summarizing correlations between cluster-level latent coordinates
       and cluster-resolved observable variables.
   - seed_42_Temporal_Latents.png
       Temporal trajectories of learned latent variables across simulation time.

Interpretation
--------------
This archive should be interpreted as supporting material for a representative
simulation run and its associated latent-space analyses. It is not intended to
imply exact deterministic reproducibility of every autoencoder-derived quantity
across retrainings, but rather to document the specific files associated with
the reported seed_42 example.

Software context
----------------
These files were generated using the Evoscope framework and its associated
autoencoder analysis scripts, as described in the manuscript and in the
corresponding GitHub repository: https://github.com/lunanfoldomics/evoscope.git

Author
------
Luca Zammataro
Lunan Foldomics LLC
