## Supplementary material for "A Minimal Regulatory Spatial Model for Emergent Multicellular Organization in Dissipative Environments": Evoscope-SI-2026.04.24.pdf

Cross-seed consistency of latent–observable relationships in Evoscope

Luca Zammataro
Lunan Foldomics LLC, Houston, TX, USA


April 25, 2026

#### Overview

These supplementary data assess the robustness of the Evoscope framework across independent simulations. To this end, we examined 10 randomly selected seeds from the full set of valid simulation runs used in the study. For each seed, we report correlation heatmaps linking learned latent variables to global and cluster-resolved observable variables. The correlation heatmaps show that the qualitative latent–observable relationship remains stable across runs, despite possible permutation, rotation, or redistribution of individual latent axes.

#### Seed selection

Ten independent seeds were randomly selected without replacement from the full set of valid simulations used in the study. Selection was performed with a fixed randomization seed to ensure reproducibility.
Selected seeds: [**38, 40, 53, 65, 89, 90, 96, 101, 104, 107**]

#### Interpretation of cross-seed variability

The purpose of these supplementary figures is not to establish one-to-one correspondence between individual latent coordinates across independent runs. Because latent axes learned by autoencoders are not expected to align identically from seed to seed, the relevant question is whether similar qualitative latent–observable relationships recur across independent simulations. The figures below support this interpretation through consistent correlation structure across independent runs.

### Supplementary Figures: Cross-seed correlation heatmaps

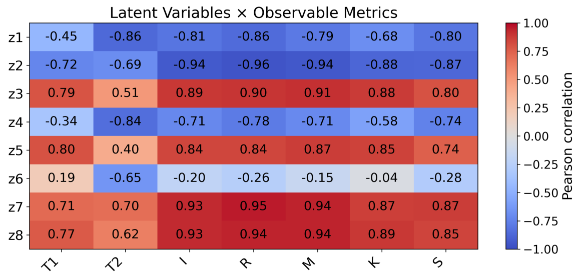


(a) Global latent vs. global observables

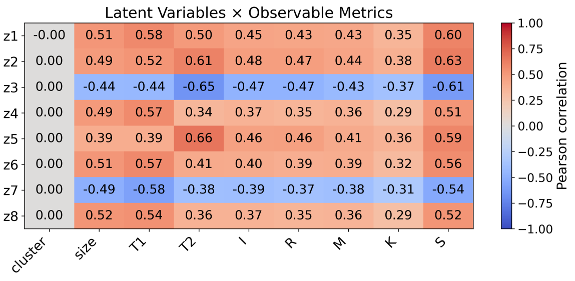


(b) Cluster-resolved latent vs. cluster observables

Figure 1: Correlation structure of the learned latent spaces for independent simulation seed 38. (A) Pearson correlations between global latent coordinates and system-wide observable variables. (B) Pearson correlations between cluster-resolved latent coordinates and cluster-level observables. Although individual latent axes are not expected to align identically across runs, the overall structure of latent–observable relationships remains qualitatively similar across independent seeds.

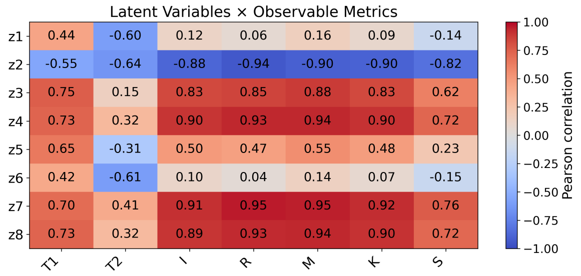


(a) Global latent vs. global observables

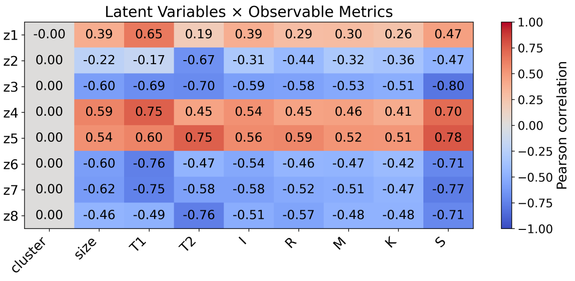


(b) Cluster-resolved latent vs. cluster observables

Figure 2: Same as Figure 1, for independent simulation seed 40.

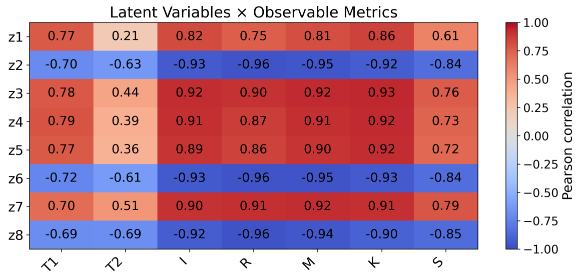


(a) Global latent vs. global observables

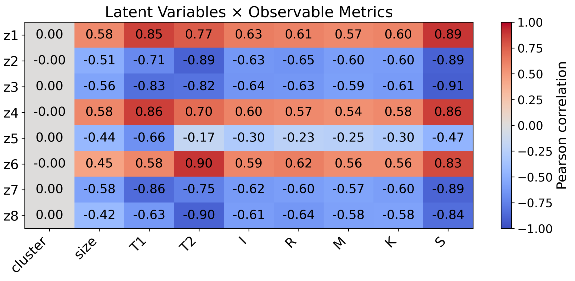


(b) Cluster-resolved latent vs. cluster observables

Figure 3: Same as Figure 1, for independent simulation seed 53.

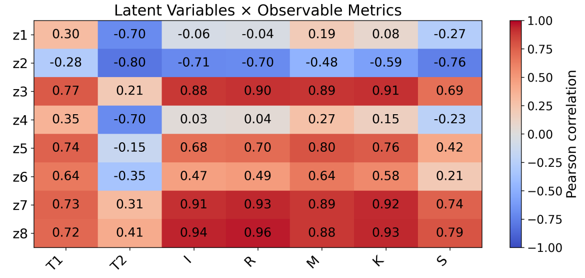


(a) Global latent vs. global observables

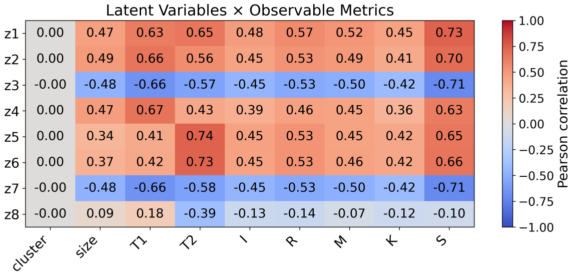


(b) Cluster-resolved latent vs. cluster observables

Figure 4: Same as Figure 1, for independent simulation seed 65.

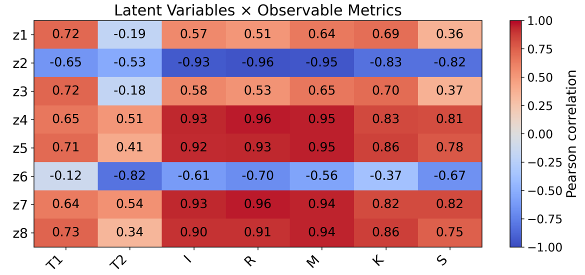


(a) Global latent vs. global observables

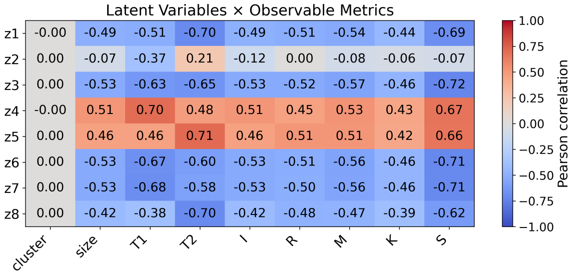


(b) Cluster-resolved latent vs. cluster observables

Figure 5: Same as Figure 1, for independent simulation seed 89.

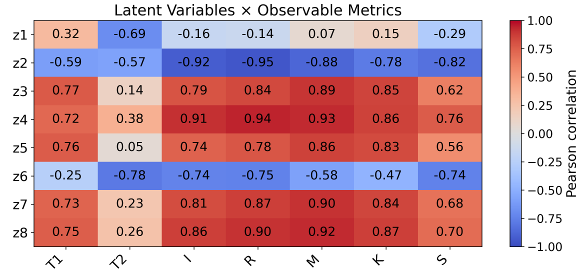


(a) Global latent vs. global observables

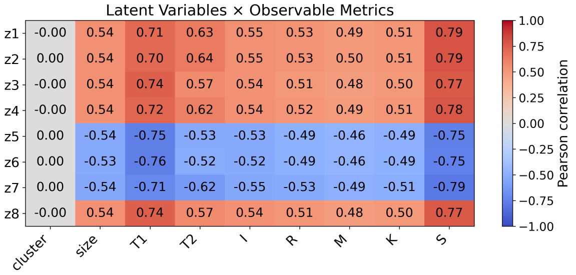


(b) Cluster-resolved latent vs. cluster observables

Figure 6: Same as Figure 1, for independent simulation seed 90.

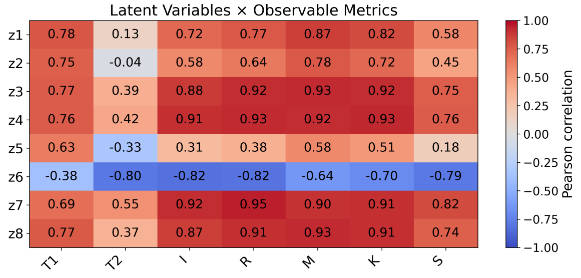


(a) Global latent vs. global observables

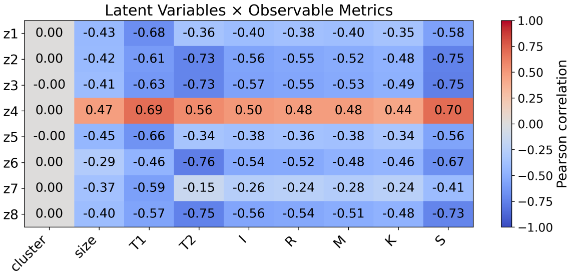


(b) Cluster-resolved latent vs. cluster observables

Figure 7: Same as Figure 1, for independent simulation seed 96.

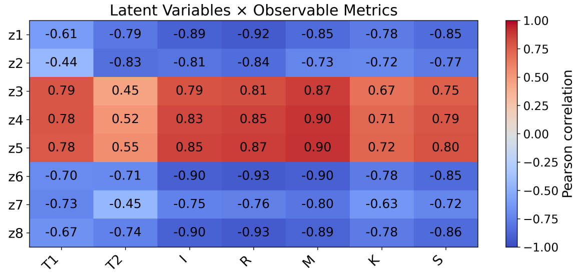


(a) Global latent vs. global observables

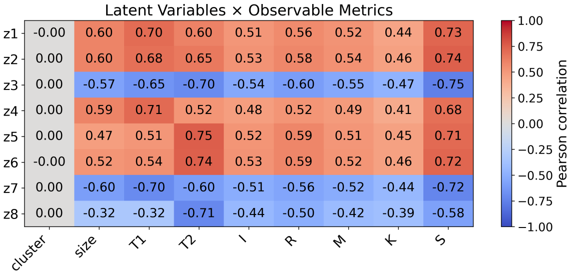


(b) Cluster-resolved latent vs. cluster observables

Figure 8: Same as Figure 1, for independent simulation seed 101.

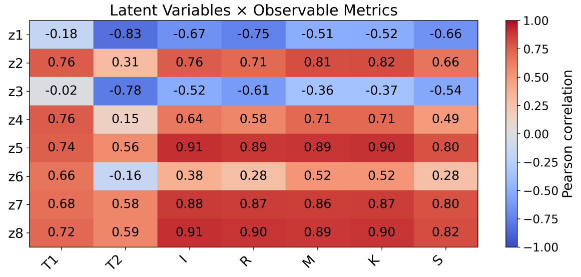


(a) Global latent vs. global observables

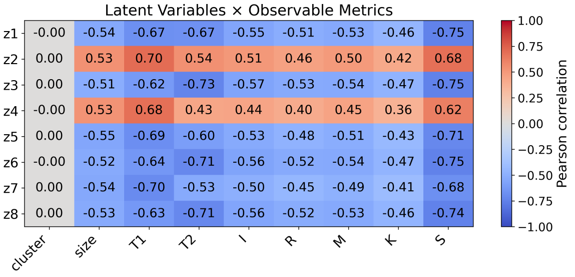


(b) Cluster-resolved latent vs. cluster observables

Figure 9: Same as Figure 1, for independent simulation seed 104.

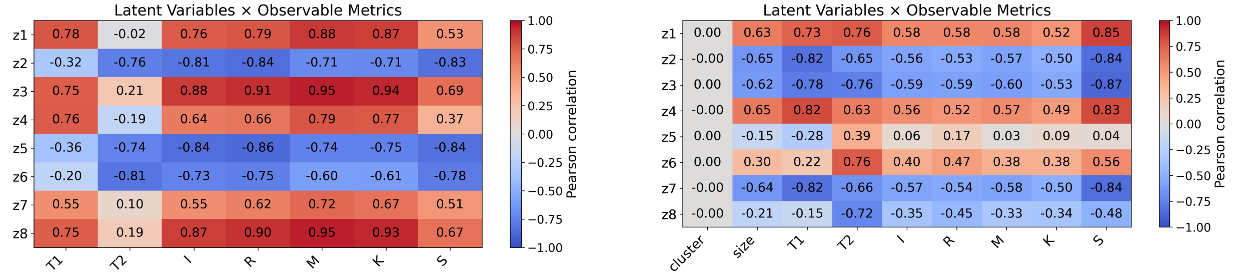


(a) Global latent vs. global observables

(b) Cluster-resolved latent vs. cluster observables

Figure 10: Same as Figure 1, for independent simulation seed 107.

#### Interpretive conclusion

The variability observed across independent heatmaps does not imply absence of latent mesoscopic structure. In representation-learning settings, independently trained autoencoders are not expected to recover identical latent axes, since equivalent internal organization may be expressed through rotation, permutation, sign inversion, or redistribution of information across coordinates. The relevant question is therefore not strict axis-by-axis reproducibility, but whether related latent–observable association patterns recur across independent runs. In this sense, the present supplementary analysis should be interpreted as evidence that Evoscope supports learnable and partially robust mesoscopic latent structure linking emergent morphology to underlying regulatory variables, rather than as proof of a unique or fully identifiable latent coordinate system.
