## Supplementary figures and images for "A Minimal Regulatory Spatial Model for Emergent Multicellular Organization in Dissipative Environments"

### seed_42_cluster_latent_heatmap.png

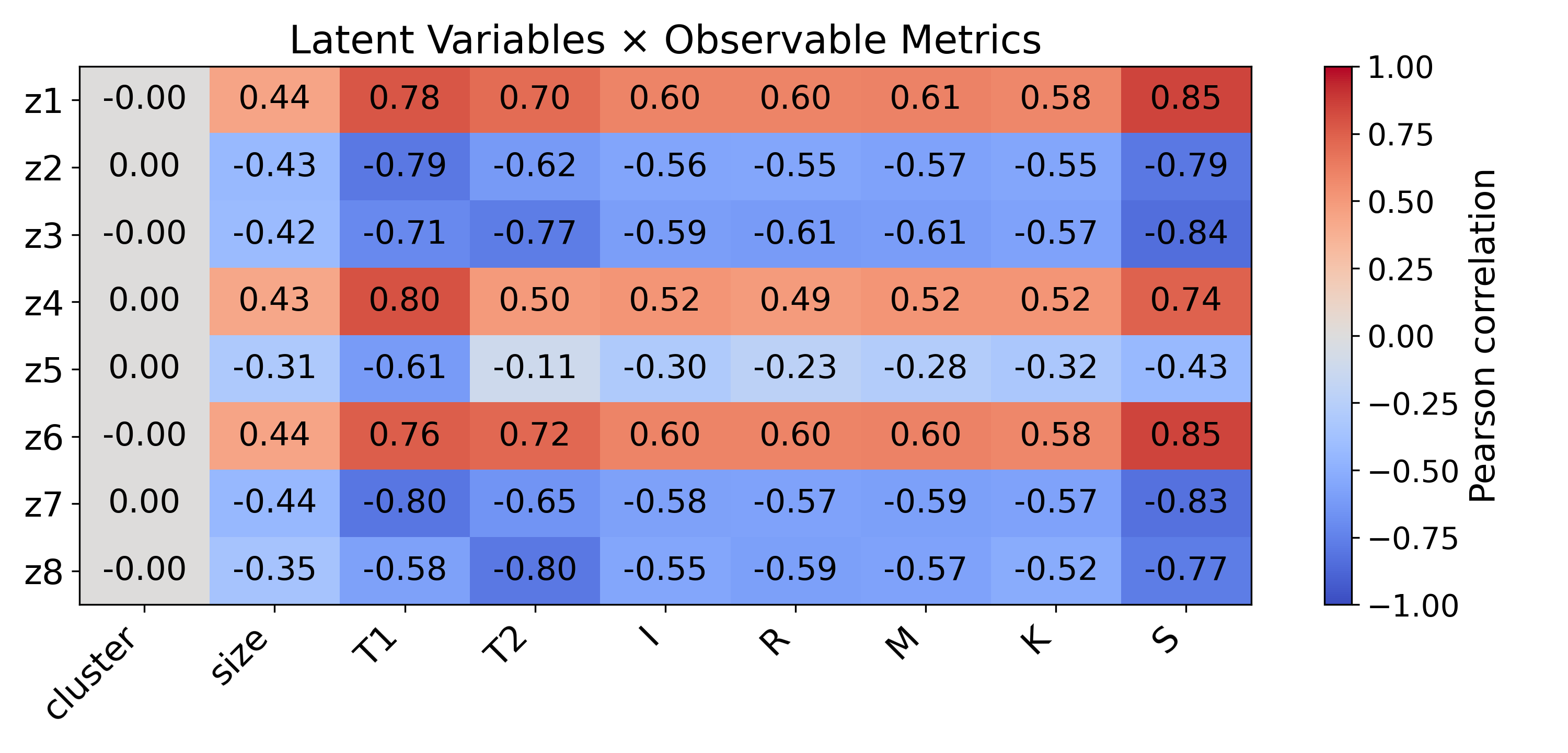

### seed_42_global_latent_heatmap.png

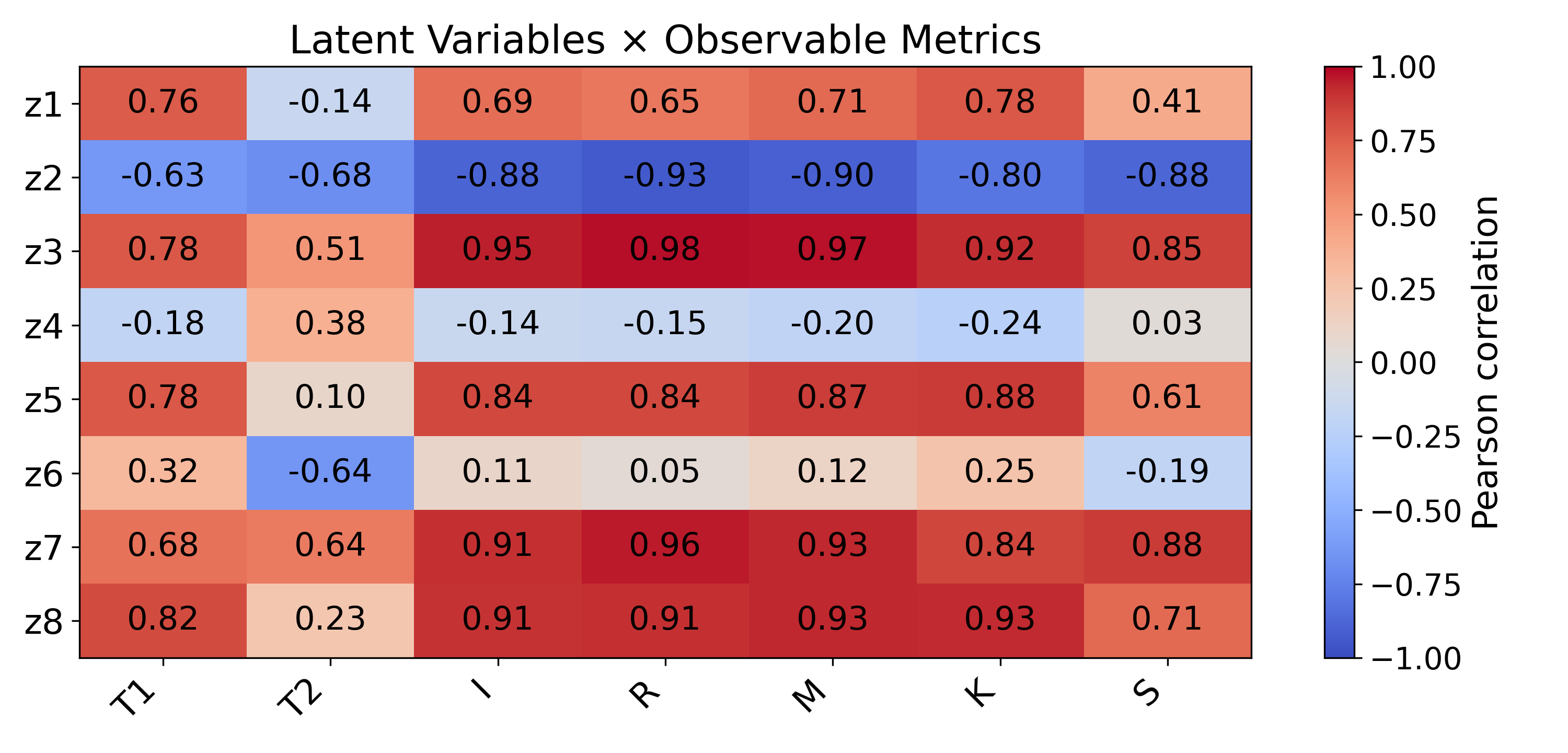

### seed_42_Temporal_Latents.png

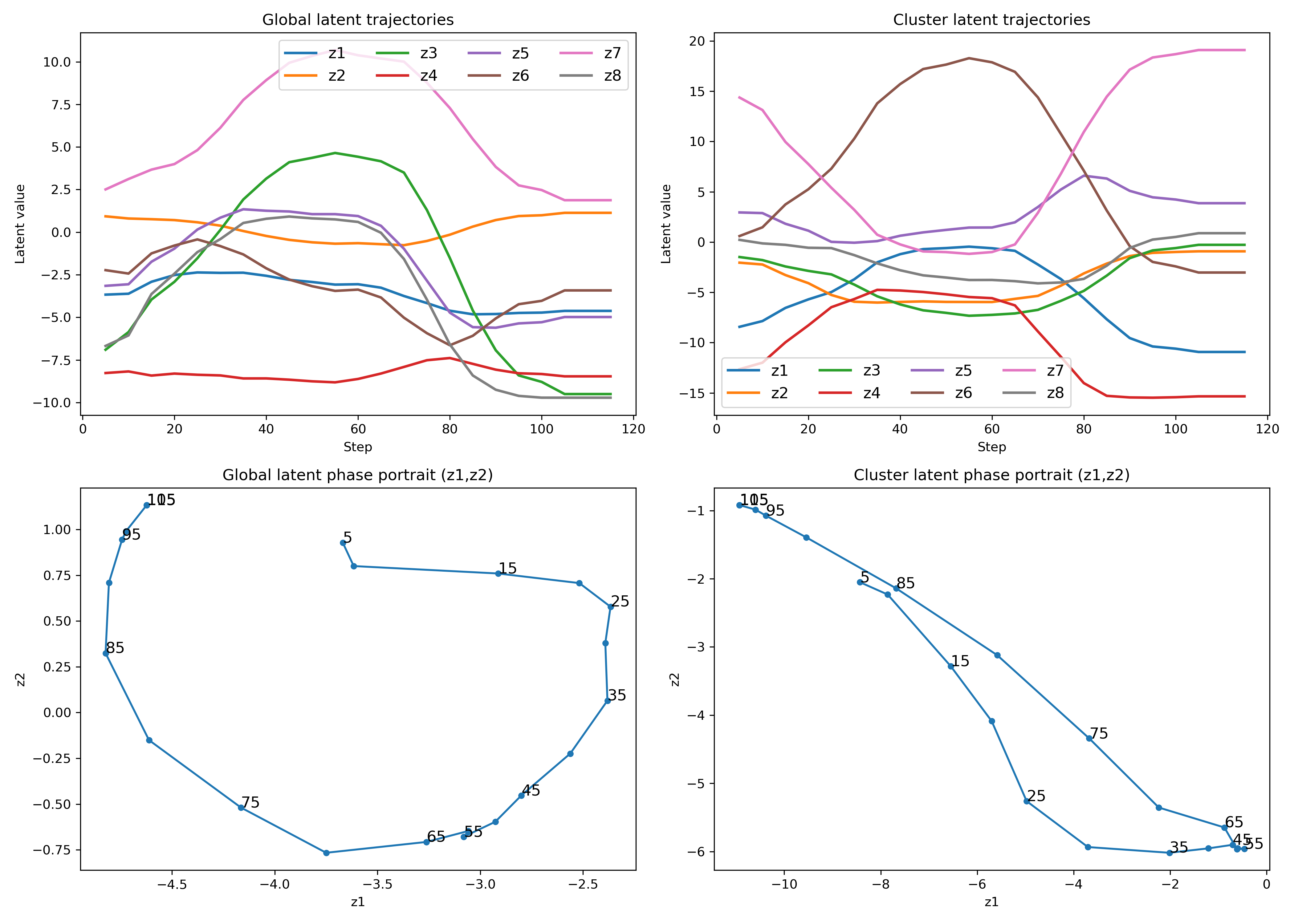
